# Endogenization and excision of human herpesvirus 6 in human genomes

**DOI:** 10.1101/2019.12.19.882522

**Authors:** Xiaoxi Liu, Shunichi Kosugi, Rie Koide, Yoshiki Kawamura, Jumpei Ito, Hiroki Miura, Nana Matoba, Motomichi Matsuzaki, Masashi Fujita, Anselmo Jiro Kamada, Hidewaki Nakagawa, Gen Tamiya, Koichi Matsuda, Yoshinori Murakami, Michiaki Kubo, Kei Sato, Yukihide Momozawa, Jun Ohashi, Chikashi Terao, Tetsushi Yoshikawa, Nicholas F. Parrish, Yoichiro Kamatani

## Abstract

The genome of human herpesvirus 6 (HHV-6) is integrated within the nuclear genome of about 1% of humans, but how this came about is not clear. HHV-6 integrates into telomeres, and this has recently been associated with polymorphisms affecting *MOV10L1*. *MOV10L1* is located on the subtelomere of chromosome 22q (chr22q) and is required to make PIWI-interacting RNAs (piRNAs). piRNAs block integration of transposons in the germline, so piRNA-mediated repression of HHV-6 integration has been suspected. Whether integrated HHV-6 can reactive into an infectious virus is also uncertain. *In vitro*, recombination of the viral genome along its terminal direct repeats (DRs) leads to excision from the telomere and viral reactivation, but the expected single DR “scar” has not been described *in vivo*. We analyzed whole-genome sequencing (WGS) data from 13,040 subjects, including 7,485 from Japan. We found an association between integrated HHV-6 and polymorphisms on chr22q in Japanese subjects. However, association with the reported *MOV10L1* polymorphism was driven by physical linkage to a single ancient endogenous HHV-6A variant integrated into the telomere of chr22q in East Asians. We resolved the junction of the human chromosome with this viral genome using long read sequencing. Unexpectedly, an HHV-6B variant has also endogenized in chr22q; two endogenous HHV-6 variants at this locus thus account for 72% of all integrated HHV-6 in Japan. We also report human genomes carrying only one portion of the HHV-6B genome, a single DR, supporting *in vivo* excision and viral reactivation. Using WGS data from North American families, we show that the incidence of HHV-6 integration into the germline is lower than its prevalence, and that integrated HHV-6 is not associated with the reported variant in *MOV10L1*. Together these results explain the recently reported association between integrated HHV-6 and *MOV10L1/*piRNAs, suggest exaptation of HHV-6 in its coevolution with human chr22q, and clarify the evolution and risk of reactivation of the only intact non-retroviral genome known to be present in human germlines.

SIGNIFICANCE STATEMENT

Human herpesvirus 6 (HHV-6) infects most people during childhood, usually only causing fever and rash. Reactivation of HHV-6 has been linked to a number of neurological diseases including encephalitis, Alzheimer’s disease and multiple sclerosis. However, about 1% of people are born with the HHV-6 genome present within their genome, included in the end “cap” of one of their 46 chromosomes. Little is known about how and when HHV-6 genomes entered human genomes, whether or not they still do, and whether or not this poses risk for virus reactivation. We looked for HHV-6 in genome sequences from over 13,000 people. Most HHV-6 variants present in human genomes have been co-evolving with human chromosomes for many generations, and new integration events are rare. Surprisingly, in almost three fourths of Japanese people with HHV-6 in their genome, HHV-6 integrated in the same end of the same chromosome – 22q. Persistence of the HHV-6 genome within the short “cap” that preserves the end of chromosome 22q suggests that the integrated viral sequence may have taken on a useful function for this chromosome. We also found that some human genomes harbor only one part of the HHV-6 genome. This part is the same part that remains after experimental viral reactivation, during which most of the virus is cut out of the genome. This warrants assessment of the risk that integration of HHV-6 into inherited human genomes is not irreversible, and possibly leads to production of infectious virus.

## INTRODUCTION

HHV-6 are *betaherpesviruses* and consist of two recently distinguished species, HHV-6A and HHV-6B, whose genomes share 90% nucleotide identity (Ablashi et al., 2014). HHV-6 are members of the *roseolavirus* family, named after roseola, the clinical syndrome of fever and rash caused by primary infection by these viruses. Most people are infected with HHV-6 as infants. In Japan, North America, and the United Kingdom, HHV-6B is most often responsible for primary HHV-6 infection. While often benign, primary HHV-6 infection can lead to central nervous system disease including febrile status epilepticus {Epstein, 2012}. Like other herpesviruses, HHV-6 can establish presumably life-long, latent infection. In contrast to other herpesviruses, this may require integration of the viral genome into the host chromosome {Arbuckle, 2010}. HHV-6 viremia occurs in about 40% of immunosuppressed transplant recipients, in whom it can cause severe complications, including limbic encephalitis {Ongradi, 2017}. HHV-6 have also been associated with other neurological diseases including multiple sclerosis (reviewed in {Leibovitch, 2014}) and Alzheimer’s disease {Readhead, 2018}.

Over 20 years ago, both representatives of HHV-6 were shown to have integrated into human chromosomes *in vivo* and been transmitted via the germline {Daibata, 1998}{Daibata, 1999}. These early reports were controversial, because it was unclear whether the viral genome was itself inherited, linked to the human chromosome, or if a human chromosomal variant that allowed the virus to integrate into a common integration site in somatic cells was inherited {Luppi, 1998}{Torelli, 1995}. More recently, both HHV-6 species have been shown to integrate into human chromosomes *in vitro* {Arbuckle, 2013}. HHV-6 integrates specifically into telomeres{Arbuckle, 2010}. The viral genome consists of terminal direct repeats (DR_L_ and DR_R_) flanking a unique region, which encodes most proteins. Each DR contains two stretches of the telomere hexameric repeat (TTAGGG)_n_, which are important for integration *in vitro* {Wallaschek, 2016A}. While homologous recombination has thus been proposed to be involved, the exact mechanisms responsible for viral integration are not clear {Wallaschek, 2016B}{Wight, 2018}.

Chromosomally-integrated HHV-6 can recombine along the DRs *in vitro*, leading to excision of the majority of the viral genome and production of infectious virus {Borenstein, 2009}{Huang, 2014}{Prusty, 2013}. This excision can leave a single DR “scar” remaining in the human chromosome. *In vivo*, one subject with X-linked SCID has been described to have become viremic with infectious HHV-6A due to reactivation of their germline-integrated viral genome {Endo, 2014}, and two infants were proposed to be infected *in utero* by a reactivated form of the virus integrated into their mother’s germline genome {Gravel, 2013}. In these cases, the single-DR genomic “scar” potentially resulting from excision and reactivation was not studied. One subject with apparent integration of a single DR in a non-telomeric location has been described {Gulve, 2017}. Two subjects with single DRs, potentially in the germline telomere, have been mentioned in discussion, but details are lacking {Huang, 2014}. It is therefore not clear how often integrated HHV-6 is excised and reactivated *in vivo*, yet this question is important to understanding the risks associated with this condition {Gravel, 2015}.

Many questions about how the HHV-6 genome enters and exits human chromosomes remain unanswered. However, with increasing WGS surveys of human populations, the prevalence of HHV-6 integration is becoming more clear. WGS of blood cells from over 8,000 subjects, mostly of European ancestry, revealed that sequences from HHV-6A or HHV-6B could be found at read depth suggesting integration into the germline genome in 0.5% of subjects {Moustafa, 2017}. Consistent with chromosomal integration, chimeric reads spanning the integration site or hybrid paired-end reads, with one end matching the virus and one end matching the human chromosome, could be found in about half of these subjects. From the globally diverse 1000 Genome Project (1kGP), WGS data from 2,535 subjects was screened and HHV-6 chromosomal integration in 0.44% on the basis of HHV-6-mapping read depth {Telford, 2018}. In the largest study addressing this topic to date, over 140,000 subjects from China were sequenced genome-wide at low depth (0.3x) using cell-free DNA collected for non-invasive prenatal testing {Liu, 2018} (hereafter, Liu *et al*.). Combining HHV-6A and HHV-6B, 0.46% had viral read depths suggestive of integration of HHV-6. Strikingly, GWAS performed on subjects with integrated HHV-6 identified a strong association with SNPs on chr22q that affect expression of a gene, *MOV10L1*, involved in production of PIWI-interacting RNAs (piRNAs). piRNAs are known to silence transposable elements (TEs), including the retrotransposons used to elongate some insect’s telomeres{Saito, 2006}{Tatsuke, 2010}. piRNAs have also been proposed to enable heritable antiviral immune memory when generated from integrated viral sequences {Parrish, 2015}{Whitfield, 2017}. Liu *et al*. interpreted the observed association to suggest that piRNAs repress HHV-6A/B integration, and that polymorphisms affecting *MOV10L1* allow for more efficient integration of HHV-6A/B during gametogenesis.

To attempt to replicate the association between the piRNA pathway and integrated HHV-6 in an independent cohort, we used WGS data from a Japanese national WGS and biobanking project, BioBank Japan (BBJ). While our GWAS identifies variants on chr22q that are highly associated with integrated HHV-6, our interpretation differs substantially from that of Liu *et al*.: GWAS signals are driven by marker SNPs in linkage disequilibrium with “founder” integrated HHV-6 variants that share ancestry, including one HHV-6A variant linked to the reported SNP in *MOV10L1*. We further characterize the ancestral integration site using long-read sequencing. Unexpectedly, two independent ancestral integrations of HHV-6 into chr22q are relatively prevalent, accounting for 72% of all integrated HHV-6 in the Japanese population. This observation, together with the expansion of the long HHV-6A-linked haplotype in the Japanese population, suggests possible functional coevolution of HHV-6 with this chromosome arm/telomere. We also describe molecular evidence of the recombination event that has been proposed to lead to HHV-6 reactivation from the integrated form, but lack *in vivo* reports. Using WGS data from families, we estimate an upper limit for ongoing HHV-6 integration into germlines. In sum, we leverage human genome sequencing to address both new and long-standing, clinically relevant questions about chromosomal integration of HHV-6; the unexpected answers raise new hypotheses about these viruses’ coevolution with humans.

## RESULTS

### Endogenization of HHV-6A on chr22q in East Asians

We first analyzed WGS data from a total of 7,485 Japanese individuals from BBJ. 3,256 subjects were sequenced to achieve high read depth (15–35×), and 4,229 subjects were sequenced at intermediate-depth (3.5×). For screening purposes, we mapped reads that failed to map to the human reference genome (hg19) to an HHV-6A reference genome. We then calculated the depth of HHV-6 genome coverage for each subject. We set a depth of 30% relative to the autosomes as a threshold consistent with chromosomal integration. We selected 32 subjects meeting this threshold as those who would have been considered to have integrated HHV-6 based on similar studies (Figure 1). None of these subjects were closer than fourth-degree relatives to each other (see methods). This suggests a prevalence of integrated HHV-6 in Japan of 0.43%. Consistent with previous results, we detected hybrid virus/human paired-end reads in 12 of these subjects {Moustafa, 2017}. However, the human chromosome-derived reads from these mate pairs did not map uniquely to the reference genome {Linardopoulou, 2005}. This precluded us from definitively assigning the site of integration using this data.

**Figure 1.**
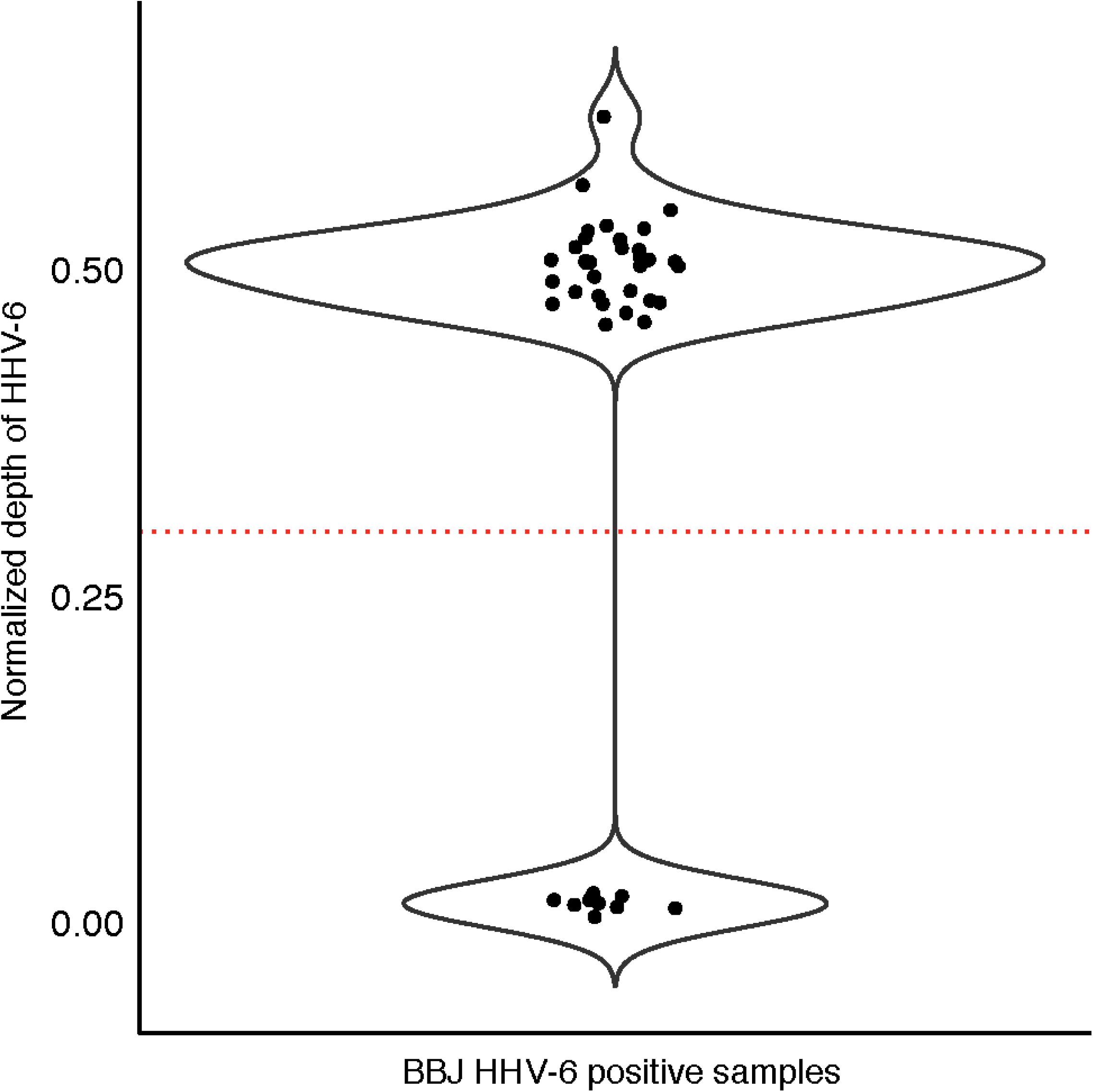
Screening for integrated HHV-6 in subjects from Biobank Japan. Unmapped WGS reads were mapped to HHV-6A (reference genome U1102). Each dot represents the depth of coverage of HHV-6 for a given subject normalized by the WGS depth of that subject. Individuals with normalized depth greater than 0.3 (dashed line) were inferred to carry integrated HHV-6 (N = 32). There is a second cluster consisting of samples with low depth of coverage of HHV-6 (N =9).

We first attempted to replicate the GWAS result reported by Liu *et al*. Despite performing GWAS with only 32 case subjects, we also identified variants at the distal end of chr22q that are highly associated with integrated HHV-6A/B (Figure 2). The previously reported index SNP rs73185306 was modestly associated with HHV-6A/B (P = 0.013, OR = 2.38); the lead SNP identified in our study is closer to the chr22q telomere and in linkage disequilibrium with more centromeric variants (Table S1). The previous association study grouped subjects with HHV-6A and HHV-6B together, under the stated rationale that they co-occurred frequently and were potentially misclassified due to sequence homology. Using the 10- to 100-fold higher sequencing depth in our study, we first distinguished subjects with integrated HHV-6A from those with integrated HHV-6B. To do so, we extracted and concatenated three viral genes (U27/U43/U83, which show high divergence between viral species) from the BBJ subjects sequenced to high depth. A phylogenetic tree of these sequences along with reference HHV-6 sequences readily discriminated viral species. Furthermore, it showed that Japanese HHV-6A sequences were monophyletic (Figure 3A). In fact, across the 143,199 bp of non-repetitive viral genome sequence called from WGS data for all four subjects with integrated HHV-6A, there were only two unique nucleotide substitutions. Notably, U27/U43/U83 sequences from HHV-6A sequences from our study are also identical to those from integrated HHV-6A in the genome of a previously-sequenced Chinese subject (HG00657) {Telford, 2018}, and distinct from circulating or integrated HHV-6 sequences derived from subjects in other locations (Figure 3A). Some, but not all, of the HHV-6B sequences were also identical across the three genes concatenated for this tree (Figure 3A). To distinguish viral species in subjects with lower sequencing depth, we calculated the ratio of the variants present in the integrated virus relative to each species’ reference genome, reasoning that there would be fewer variants called for the species from which the integrated virus belonged. Plotting these values along with those of the deeply sequenced subjects showed that, even in subjects with 3.5× sequencing depth, the viral species could readily be distinguished (HHV-6A N=12, HHV-6B N=20; Figure 3B). Furthermore, no subjects appeared to harbor sequences from both HHV-6A and HHV-6B (Figure 3B).

**Figure 2.**
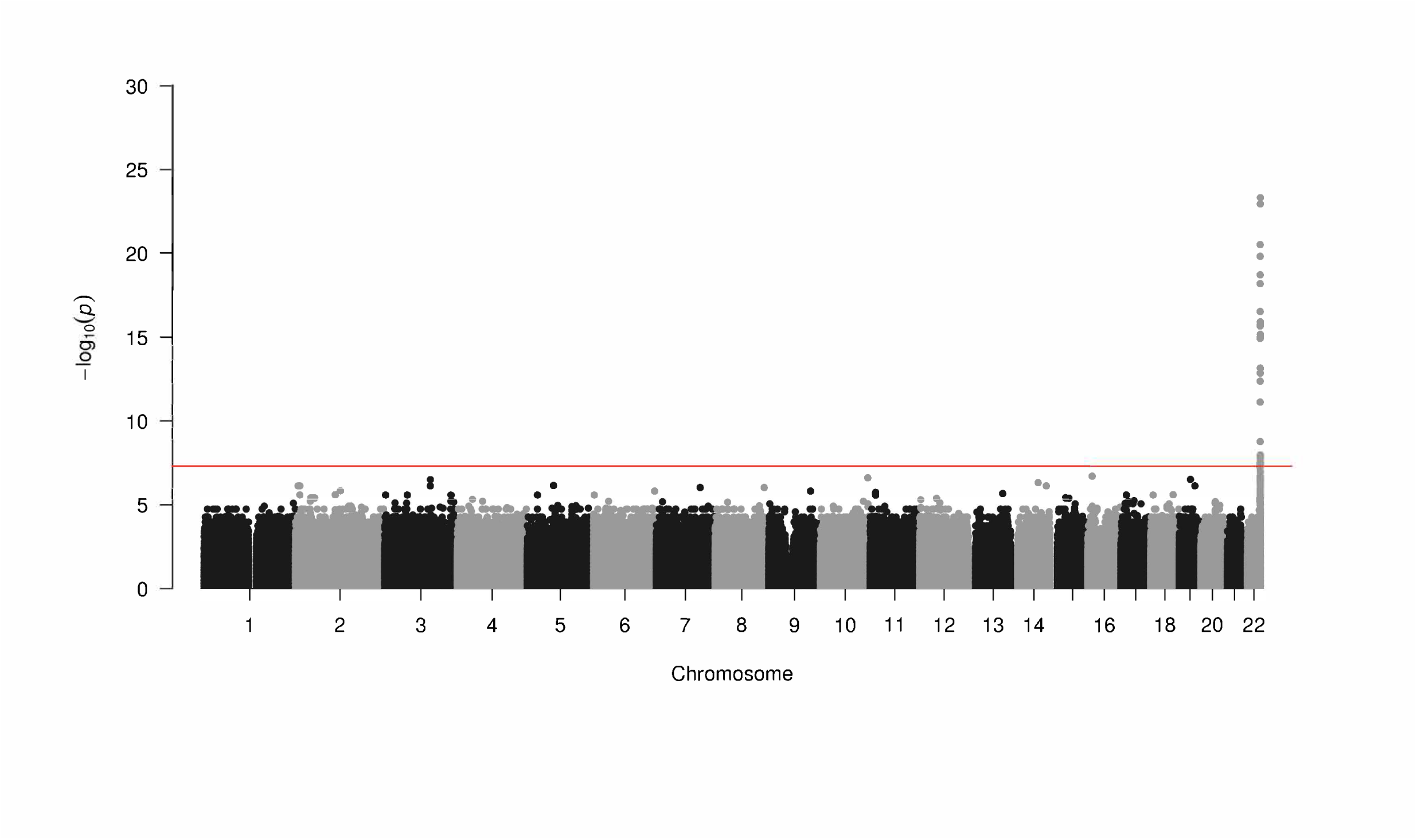
Integrated HHV-6A/B is associated with variants on chr22q. Manhattan plot presenting the P values for association between a variant and integrated HHV-6 (N = 32) comparerd to subjects who do not carry integrated HHV-6. The −log10 P value (Fisher exact test) from variants is plotted according to their physical position on successive chromosomes.

**Figure 3.**
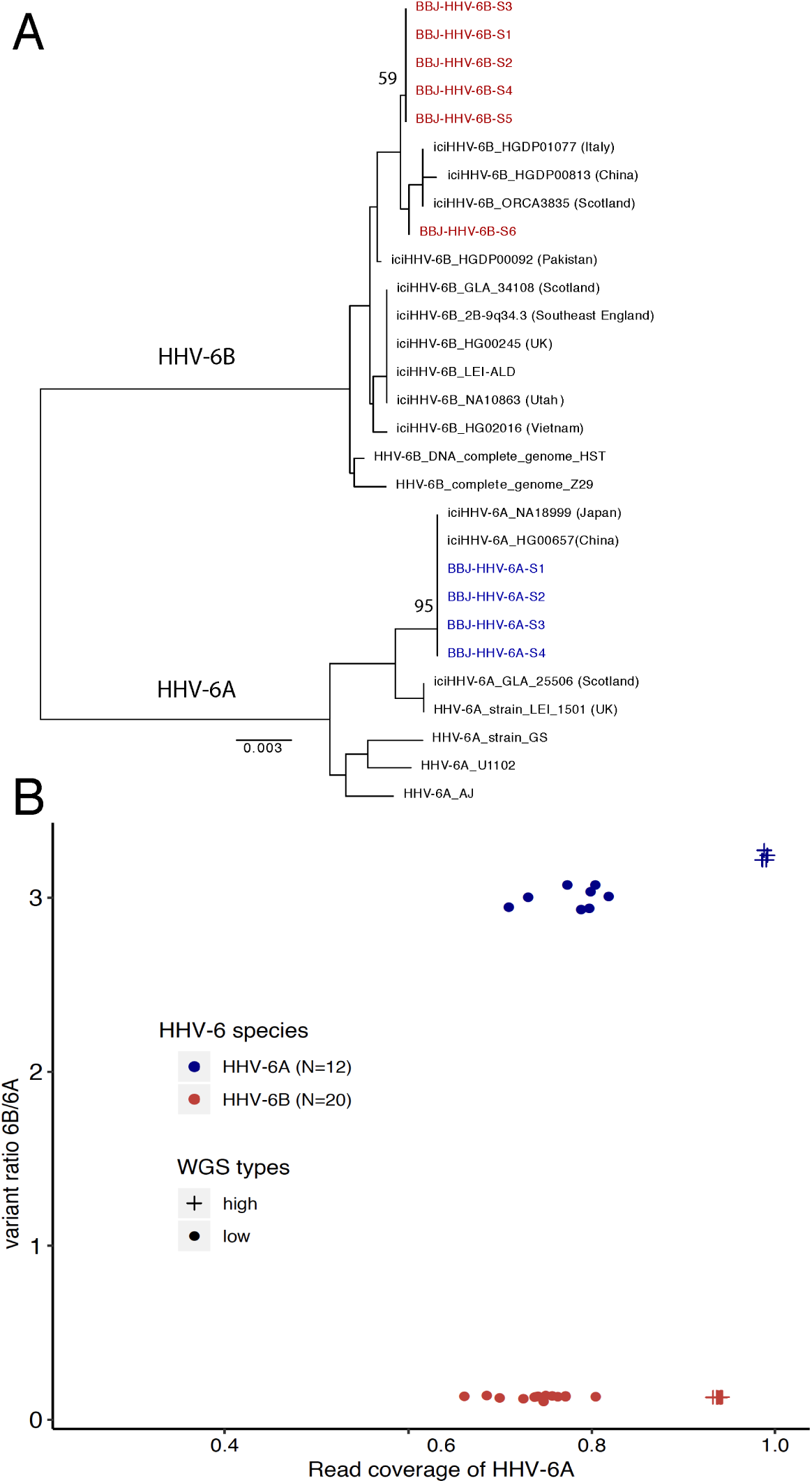
HHV-6 species is readily distinguished in subjects with both high- and low-depth WGS. **A) Neighbor-joining phylogenetic tree of concatenated HHV-6 viral genes U27/U43/U87 from 29 HHV-6 genomes.** Phylogenetic tree analysis demonstrated HHV-6A sequences from high-depth WGS samples from BioBank Japan (BBJ), colored in blue, are monophyletic and are identical to integrated HHV-6A sequences of Japanese (NA18999) and Chinese (HG00657) subjects from 1KGP. HHV-6B sequences obtained from BBJ subjects are labelled in red. Bootstrap value per 100 replicates of selected nodes is shown. The scale bar represents 0.003 substitutions per site. **B) Comparing variants and mapping coverage relative to HHV-6A and HHV-6B reference genomes distinguishes species for subjects with low-depth WGS.** The Y axis indicates the ratio of variants called in comparison to the HHV-6B reference versus those called in comparison to the HHV-6A reference genome. The X axis represents the ratio of percentage of coverage of the HHV-6B reference versus coverage of the HHV-6A reference genome. Samples determined as HHV-6A (N = 12) and HHV-6B (N = 20) are colored in blue and red respectively. Subjects sequenced in high-depth or low-depth WGS are represented by crosses and dots, respectively.

The near-identity of integrated HHV-6A viral sequences in our dataset suggested that they descended from a single integration event that increased in proportion in the population via vertical transmission. In such a scenario, the SNPs associated with integrated HHV-6 variants that are identical by descent, representing a single historical event, would not necessarily be indicative of the biological factors contributing to HHV-6 integration as initially interpreted. Instead, these SNPs may be in linkage disequilibrium with the chromosomal integration site of the ancestral integrated HHV-6A. To test this, we repeated GWAS using only subjects with integrated HHV-6A (Figure 4A). Again, despite using only 12 case subjects, GWAS revealed a highly significant association with SNPs on chr22q, including with SNPs overlapping the locus identified by Liu *et al*. (Table S2, Figure 4B). Notably, rs73185306 is significantly associated HHV-6A (P = 1.85E-05, OR = 7.36) but not HHV-6B (P = 0.578, OR = 0.54). From these results, we hypothesized that there was an ancient “founder” integration of HHV-6A that remains present in the telomere of chr22q in some East Asians and may have contributed to the association with *MOV10L1* in the previous study.

**Figure 4.**
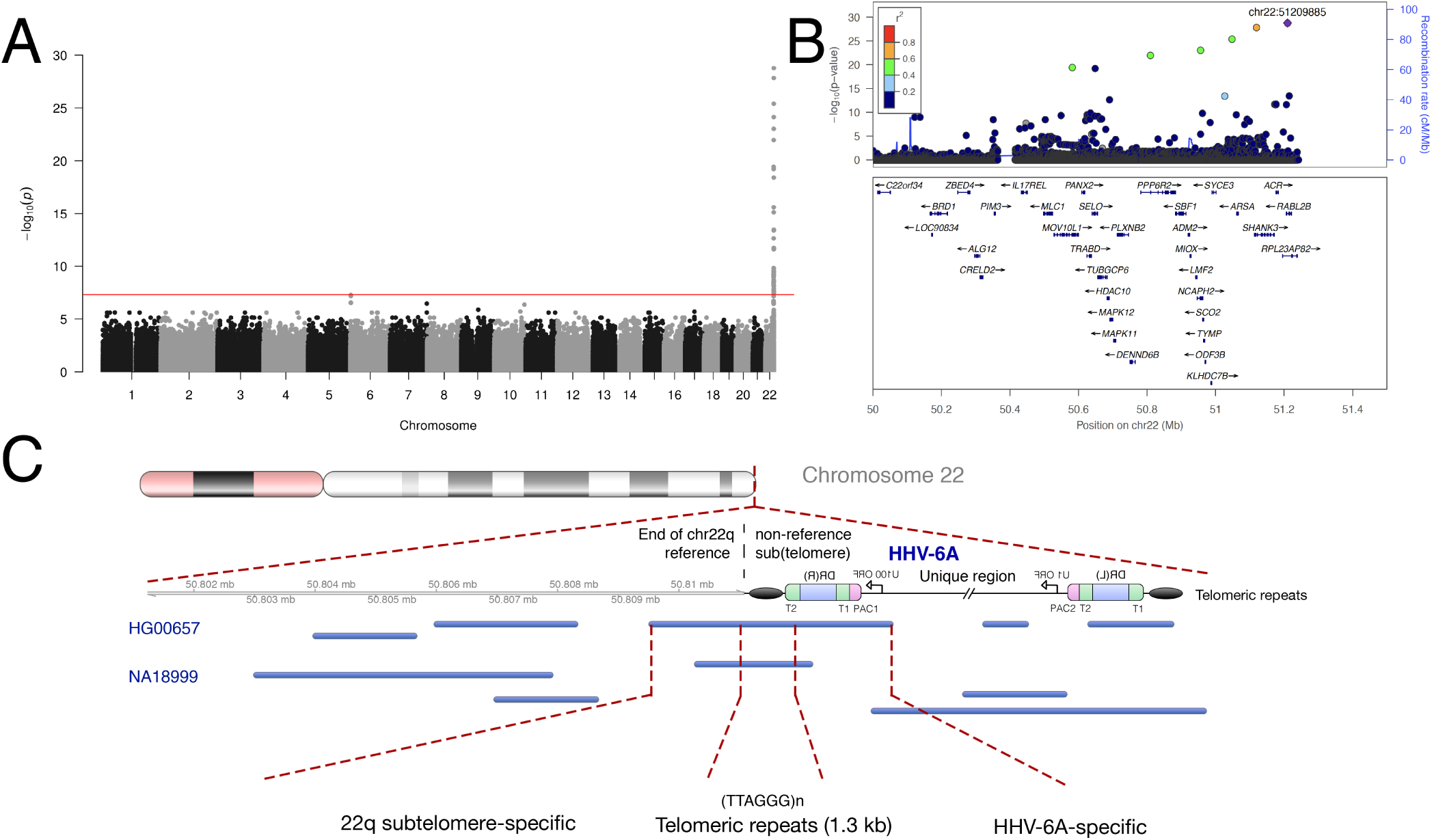
An endogenous HHV-6A variant in East Asians integrated into chr22q. **A) Manhattan plot presenting the P values for association between variant and integrated HHV-6A (N = 12).** The −log10 P (Fisher exact test) from variants is plotted according to its physical position on successive chromosomes. **B) Regional association plot of the 22q region.** The −log10 P (Fisher exact test) for association in the GWAS of iciHHV-6A are shown. Proxies are indicated with colors determined from their pairwise r2 from the high-depth BBJ WGS data (red, r2 > 0.8; orange, r2 = 0.5 −0.8; yellow, r2 = 0.2–0.5; white, r2 < 0.2 or no information available). **C) Long-read sequencing identifies endogenous HHV-6A integration site.** Mapping of individual long sequencing reads (black lines) to the chr22q reference sequence and to HHV-6A is depicted. Reads were obtained from lymphoblastoid cell lines derived from two subjects with integrated HHV-6A (HG00657 and NA18999) who bear the rare variant rs566665421. The reads that span the integration site are highlighted, demonstrating the integration site of HHV-6A in both subjects is the non-reference terminal heterochromatin of q the arm of chr22.

The lead SNP from Liu *et al*. is 780 kb centromeric to the chr22q telomere, into which we suspected an ancestral integration of HHV-6A. We thus hypothesized the existence of an extended haplotype comprising at least the last ∼780 kb of 22q, spanning from the telomere to this SNP. To test for such a haplotype, we used phase-estimated microarray data from chr22q to build a phylogenetic tree{Stephens, 2005}. This revealed that 9 subjects with HHV-6A from BBJ clustered together with a well-supported node, consistent with them sharing a haplotype on distal chr22q (Figures S1, S2). We confirmed that none of the subjects with HHV-6A from BBJ were closely related, indicating that haplotype sharing was localized at chr22q. We next tabulated rare variants highly associated with HHV-6A for subjects sequenced to high depth (Table 1). These results are concordant with the haplotype tree; presence of the most telomeric rare variant associates perfectly with integrated HHV-6A. Some subjects carry the centromeric but not the telomeric rare variants, suggesting a recombination event, and these subjects lack integrated HHV-6A. Subject HG00657 lacks most of the rare variants present on the shared haplotype, explaining why this subject did not cluster with Japanese subjects with integrated HHV-6A (Figure S1). However, this subject shares the most distal rare variant (rs566665421) associated with HHV-6A. This, along with the near identity of the viral sequences, suggests that an integrated HHV-6A variant present in both China and Japan is the result of ancestral viral integration into chr22q.

**Table 1:** Rare variants in chromosome 22q subtelomeric region co-segregate with integrated HHV-6A

| ID | Group (2) | Position (centromeric to telomeric) (1) |  |  |  |  |  |  |  |  | HHV-6 status |
| --- | --- | --- | --- | --- | --- | --- | --- | --- | --- | --- | --- |
|  |  | 22:50522524<br>rs73185306* | 22:50582094<br>rs149078280 | 22:50810165 | 22:50874466 | 22:50874474 | 22:50956116<br>rs200663937 | 22:51048089<br>rs531808350 | 22:51058555 | 22:51209885<br>rs566665421 |  |
| HHV-6A-S1 | BBJ | C/T | C/T | C/G | G/GCGTTT | G/A | C/T | T/C | T/TA | A/G | HHV-6A |
| HHV-6A-S2 | BBJ | C/T | C/T | C/G | G/GCGTTT | G/A | C/T | T/C | T/TA | A/G | HHV-6A |
| HHV-6A-S3 | BBJ | C/T | C/T | C/G | G/GCGTTT | G/A | C/T | T/C | T/TA | A/G | HHV-6A |
| HHV-6A-S4 | BBJ | C/T | C/T | C/G | G/GCGTTT | G/A | C/T | T/C | T/TA | A/G | HHV-6A |
| NA18999 | 1KGP (Japan) | C/T | C/T | C/G | G/GCGTTT | G/A | C/T | T/C | T/TA | A/G | HHV-6A |
| HG00657 | 1KGP (China) | C/C | C/C | C/C | G/G | G/G | C/C | T/T | T/T | A/G | HHV-6A |
| CTRL(JPN)-44 | BBJ | C/T | C/T | C/G | G/GCGTTT | G/A | C/T | T/C | T/TA | A/A | No HHV-6 reads |
| CTRL(JPN)-34 | BBJ | C/T | C/T | C/C | G/G | G/G | C/C | T/T | T/T | A/A | No HHV-6 reads |
| HHV-6B-S1 | BBJ | C/C | C/C | C/C | G/G | G/G | C/C | T/T | T/T | A/A | HHV-6B |
| HHV-6B-S2 | BBJ | C/C | C/C | C/C | G/G | G/G | C/C | T/T | T/T | A/A | HHV-6B |
| HHV-6B-S3 | BBJ | C/C | C/C | C/C | G/G | G/G | C/C | T/T | T/T | A/A | HHV-6B |
| HHV-6B-S4 | BBJ | C/C | C/C | C/C | G/G | G/G | C/C | T/T | T/T | A/A | HHV-6B |
| HHV-6B-S5 | BBJ | C/T | C/C | C/C | G/G | G/G | C/C | T/T | T/T | A/A | HHV-6B |
| HHV-6B-S6 | BBJ | C/C | C/C | C/C | G/G | G/G | C/C | T/T | T/T | A/A | HHV-6B |

1) position based on reference Hg38 and dbSNP ID, if applicable
2) BBJ, Biobank Japan; 1KGP, from 1000 Genomes Project subjects with iciHHV-6 (Telford et al., 2018)
\* Liu et al. index SNP

We considered that linkage disequilibrium (LD) between this integrated HHV-6A genome and the SNP reported by Liu *et al*. might explain their association result. However, if that were the case, we reasoned that the most telomeric variant (rs566665421), rather than one in *MOV10L1*, would be most highly associated SNP with HHV-6A/B integration. We checked this site in a database providing summary statistics from Liu *et al*. (https://db.cngb.org/cmdb/), but genotyping data is not available. Therefore, we speculate that the low sequencing depth used for the previous study of integrated HHV-6 in East Asians precluded accurate genotyping of variants more closely linked to this trait, prevented recognition of a large LD block in the region of the association, and obscured evidence of shared ancestry of the integrated HHV-6A allele, resulting in identification of an index SNP distant from the “causal” HHV-6A variant present in the telomere.

To confirm that the HHV-6A-linked haplotype is physically linked to telomere-integrated HHV-6, we performed long-read sequencing. Only two subjects with genotypes publicly-available carry the rare variant rs566665421: the Chinese subject mentioned above (HG00657) and one Japanese subject (NA18999) {Genomes Project, 2015}. We hypothesized that these subjects harbor the same endogenous HHV-6A variant integrated into chr22q. We obtained lymphoblastoid cell lines derived from these subjects and performed long-read sequencing. This yielded individual reads that mapped to both HHV-6A and to the subtelomere of chr22q (Figure 4C). There are approximately 1.3 kb of hexameric repeats between the viral DR_R_, which, consistent with previous results, is oriented as the more centromeric DR in integrated HHV-6 genomes, and the terminal base of the chr22q reference sequence {Arbuckle, 2013}. To confirm physical linkage of integrated HHV-6A with chr22q in another way, we used DNA samples from 6 Japanese subjects previously identified to have inherited chromosomally integrated HHV-6A. In these subjects, the integration site had previously been mapped using fluorescent in situ hybridization (FISH) to chr22q {Miura, 2018}. Including HG00657 and NA18999, we sequenced SNPs in eight subjects with integrated HHV-6A mapped to chr22q (Table S3). The SNP reported by Liu *et al*. was present in six of these subjects, whereas all carried rs566665421. Thus both FISH mapping and direct sequencing localize an endogenous HHV-6A variant shared by Japanese and Chinese subjects to the telomere of chr22q, linked to an shared extended haplotype.

When this HHV-6A integration event occurred and when the allele entered the Japanese population may be informative about its prevalence in other populations. To estimate integration timing, we assumed that mutations arising in the HHV-6A genome after integration accumulate at the same rate as other chromosomal mutations (see methods). The number of polymorphisms present in the six deeply-sequenced HHV-6A genomes suggests that the virus integrated 30,556 years ago (95% CI 15,253-54,672) assuming a generation time of 25 years (Figure S3A). Considering polymorphisms in the HHV-6A genomes present only in Japanese subjects gives an estimate of 14,881 years (95% CI 4,832 to 34,727). Of the 11 polymorphisms observed, each observed in a single subject, 3 were in non-coding regions, 4 were missense, and 4 were synonymous. Next, we estimated how long the HHV-6A-linked haplotype has been recombining in the Japanese population. We modeled this using a simple deterministic equation based on the decay of linkage disequilibrium (LD) between integrated HHV-6A and a linked marker allele (rs149078280-T) {Nielsen, 2013}. This estimate suggested a much more recent introduction to Japan some 875 years ago (CI 250-2,350 years, Figure S3B). While divergent in estimating how recently this haplotype arrived to Japan, both models suggest that this endogenous HHV-6A allele existed in other East Asian populations prior to entering the Japanese population, perhaps during the Jomon period. Consistent with this interpretation, we observed that SNPs linked with the viral integration are present in Northeastern Asian populations at a similar frequency to these SNPs in the Japanese population (NARD Database, https://nard.macrogen.com).

### Endogenization of HHV-6B on chr22q

Subjects whose HHV-6B sequences were nearly identical (Figure S4A) also clustered on chr22q haplotype analysis (Figure S4B), unexpectedly suggesting another variant with shared ancestry. We performed GWAS using 11 subjects with this clonal integrated HHV-6B variant (Figure 5A/B). Consistent with integration into the same chromosome arm as the endogenous HHV-6A variant described above, this integrated HHV-6B variant is also associated with SNPs on chromosome 22q (Table S4). The haplotype linked to this viral genome is more common than that on which HHV-6A is integrated, with fewer significantly associated variants extending into the subtelomere (Figure 5B). To confirm that this GWAS result reflects physical linkage, we obtained DNA from three subjects previously identified to have integrated HHV-6B mapped by FISH to chr22q {Miura, 2018}. Sanger sequencing of the most highly associated variant (chr22:51184036) revealed that all were heterozygous for the minor allele, and none carried the rs73185306 variant (Table S5). Together, these results show that in the Japanese population, two different chr22q haplotypes are associated with the majority of integrated HHV-6 – one with HHV-6A and the other with HHV-6B – and that shared chr22q haplotypes correspond with shared, clonal integrated HHV-6 sequences, representing endogenous HHV-6.

**Figure 5.**
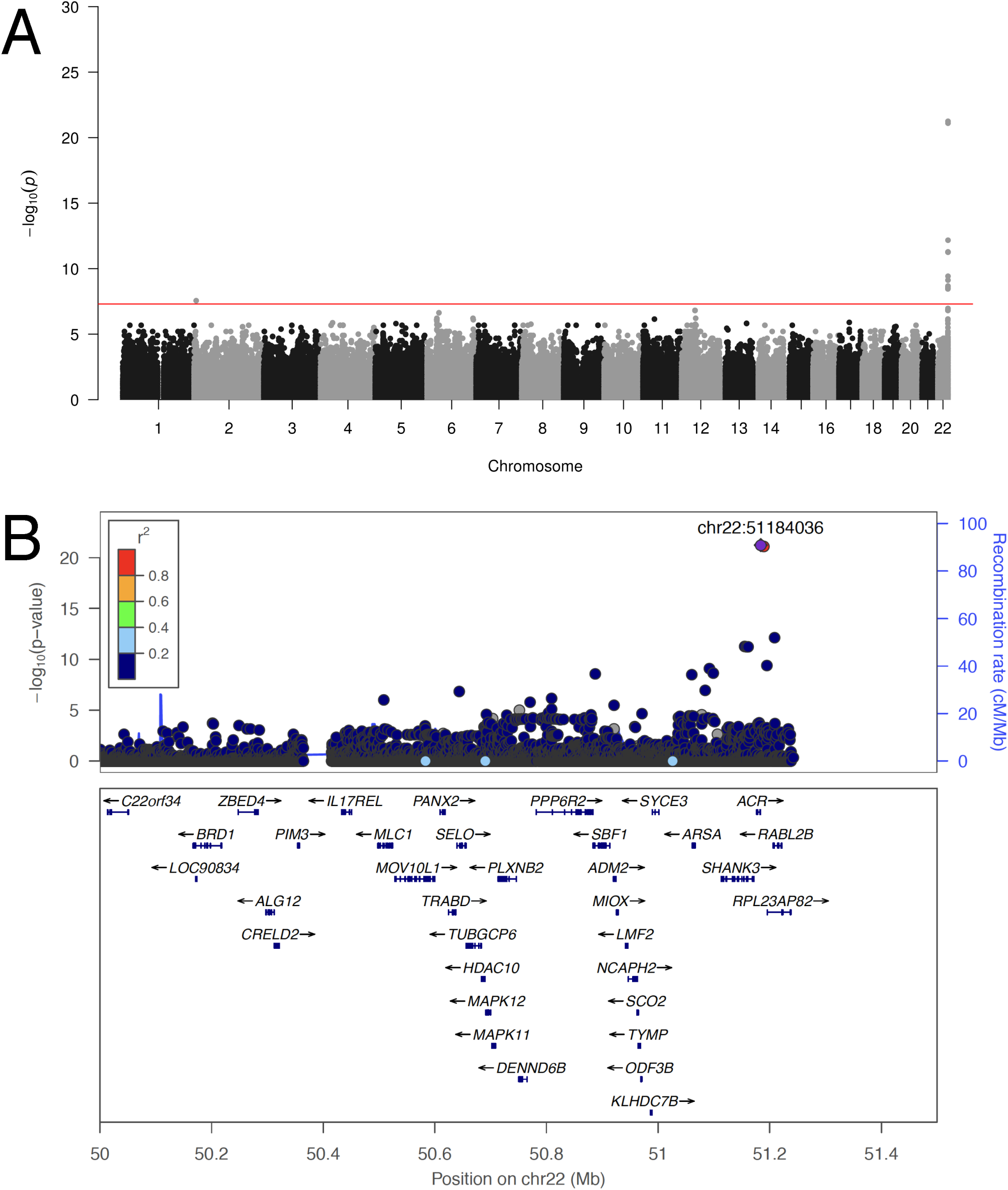
A prevalent endogenous HHV-6B variant integrated into chr22q. **A) Manhattan plot presenting the P values for association between variants and clonal integrated HHV-6B (N = 20).** The −log10 P (Fisher exact test) from variants is plotted according to its physical position on successive chromosomes. **B) Regional association plot of the 22q region.** The −log10 P (Fisher exact test) for association in the GWAS of iciHHV-6A are shown. Proxies are indicated with colors determined from their pairwise r2 from the high-depth BBJ WGS data (red, r2 > 0.8; orange, r2 = 0.5 −0.8; yellow, r2 = 0.2–0.5; white, r2 < 0.2 or no information available).

### Excision of HHV-6B from the genome

We next analyzed subjects with reads mapping to HHV-6A at a depth below the threshold used to infer germline integration. We analyzed the coverage of the viral genome in these subjects and compared to those with integrated HHV-6 described above. This revealed two distinct coverage patterns: subjects with reads mapped across the entire HHV-6 genome, and subjects with reads mapped to the DR region only (Figure S5). This partial coverage pattern was not observed in any subjects previously inferred to carry integrated HHV-6A/B. The depth of reads covering the DR region was lower in subjects who lack U region-mapped reads, suggesting that only a single DR is present in these subjects. Sequencing coverage of the DR region in these subjects terminated abruptly adjacent to the viral genome packaging sequences (Pac1/Pac2). Notably, this single DR configuration has been previously proposed as the molecular signature of recombination along the DRs, shown *in vitro* to lead to viral reactivation {Huang, 2014}.

BBJ WGS data was derived from DNA extracted from nucleated blood cells. We hypothesized that clonal expansion of a hematopoietic lineage in which recombination and excision of integrated HHV-6 had occurred could result in detection of only DR sequences from blood-derived DNA. If this were the case, the entire HHV-6 genome could be present in other cells, but at low abundance in the blood. To indirectly assess this, we obtained additional blood-derived DNA from these subjects and performed digital droplet PCR using primers for both the DR region and a well-conserved U region (Figure S6). As a control, we used DNA from subjects from whom the entire HHV-6 viral genome was detected by WGS. No evidence of low-level U-region integration, detectable by PCR but not WGS, was observed for subjects whose WGS reads mapped only to the DR region. All subjects with a single DR integration were determined to be of subtype HHV-6B using the species-specific DR probe used for PCR (Figure S6). This result argued against mosaicism as the explanation for WGS reads mapping only to the DR region.

We next performed phylogenetic analysis of integrated HHV-6B DR regions from all deeply-sequenced subjects, including four with single DR integration, to further clarify the evolution of the integrated single DRs (Figure S7). Three of the DR sequences from subjects with single DR integration were identical, and a fourth varied from these at two sites. Based on this result, we hypothesized that the single DR sequence mostly shared by these subjects could potentially reflect a single historical recombination event. To further clarify this point, we performed GWAS using 9 subjects bearing single DR HHV-6B integration (Figure 6A, B). Consistent with a single founder integration into the human chromosome and excision of the majority of the viral genome in the germline prior to vertical transmission (Figure 6C), subjects bearing single DR HHV-6B integration often shared telomere-proximal SNPs on 7q.

**Figure 6.**
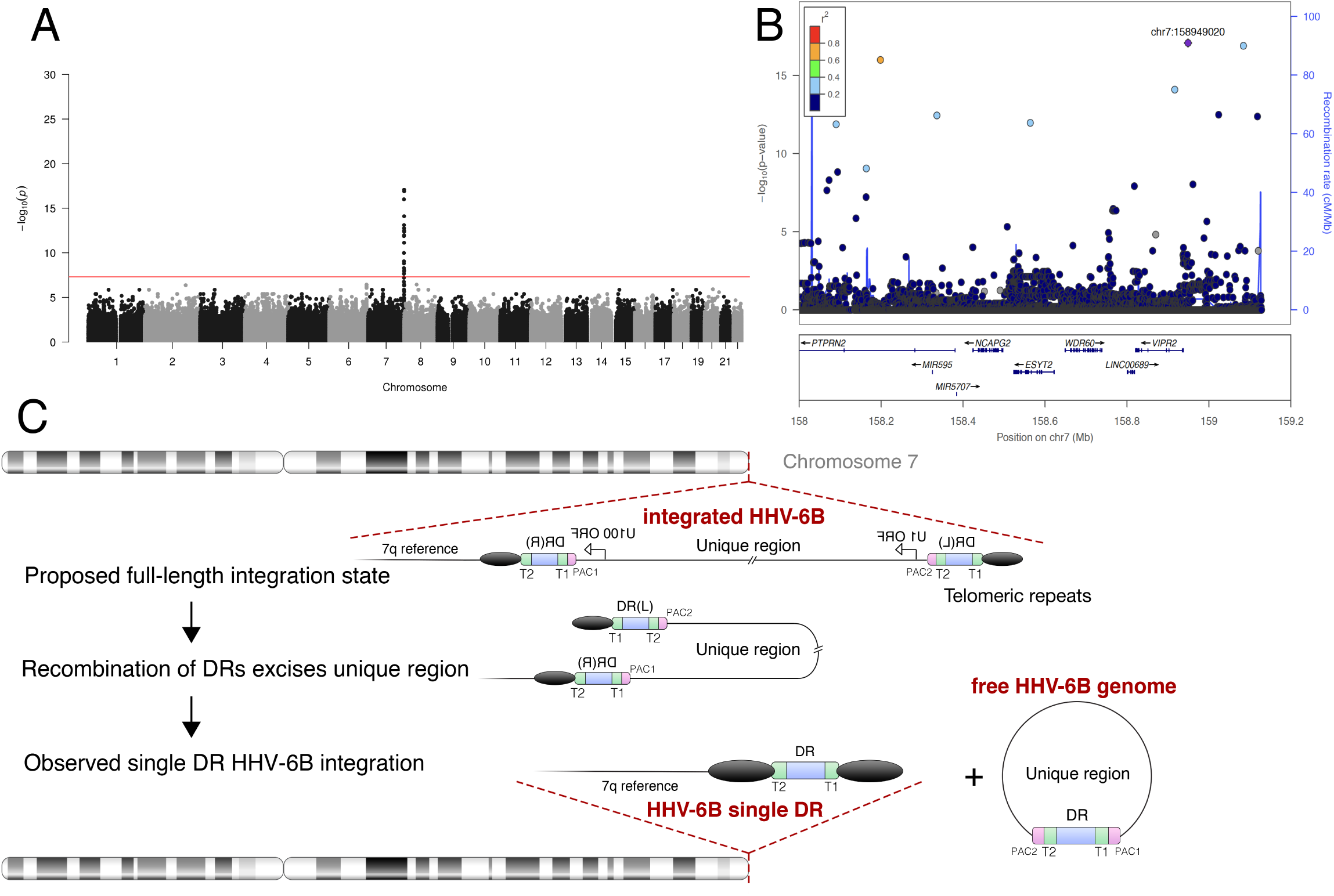
Recombination and excision of HHV-6B from chromosome 7q. **A) Manhattan plot from GWAS of subjects bearing HHV-6B single DR integration (N = 9).** The −log10 P (Fisher exact test) from variants is plotted according to its physical position on successive chromosomes. **B) Regional association plot of the 7q region.** The −log10 P (Fisher exact test) for association in the GWAS of iciHHV-6A are shown. Proxies are indicated with colors determined from their pairwise r2 from the high-depth BBJ WGS data (red, r2 > 0.8; orange, r2 = 0.5 −0.8; yellow, r2 = 0.2–0.5; white, r2 < 0.2 or no information available). **C) Model of HHV-6 recombination and excision resulting in the observed integrated single DR.** Schematic of the proposed germline recombination event (after Wood and Royle, 2017) leading to excision of the majority of integrated HHV-6B sequence resulting in the integrated single DR form.

### Incidence of HHV-6 integration

Our results to this point suggested that few integrated HHV-6 variants of shared ancestry account for majority of integrated HHV-6 in East Asians. To estimate the rate of newly incident integration of HHV-6 into the human genome, we analyzed WGS data from a total of 5,555 subjects from North American families affected by autism spectrum disorder (ASD) in the MSSNG database. We identified 1357 families with sequence data for both parents and at least one child {Jiang, 2013}. The prevalence of integrated HHV-6 among parents in the MSSNG database is 0.91%. 18 of the 30 children born to a parent with integrated HHV-6 inherited HHV-6, which is not inconsistent with the principle of independent assortment (P= 0.362 by binomial test). There was no association between integrated HHV-6 and ASD (P = 0.277). We found no Mendelian error with regards to HHV-6 integration in 1,674 children born to parents without integrated HHV-6. This suggests that the upper limit of the 95% CI for the incidence of integrated HHV-6 is 0.0018 by the “rule of 3/*n*” or 0.0023 by the Wilson score interval; thus incidence of HHV-6 integration is less a third of its prevalence in this population. This is consistent with the observation of that only two distinct endogenous HHV-6 variants account for most integrated HHV-6 in Japan. In this dataset, we found no association between integrated HHV-6A or HHV-6B with the index SNP reported by Liu *et al*., rs73185306-T (Table S6), supporting our interpretation that a specific endogenous HHV-6A variant present in East Asians is associated with *MOV10L1*.

rs73185306-T, an eQTL of *MOV10L1*, was hypothesized to influence HHV-6A/B integration by affecting piRNA function. Our data do not directly address this hypothesis. However, our results show that the hypothesis was likely proposed based on fewer independent HHV-6 integration events than initially suspected. To address the significance of this SNP on piRNA function, we tested the effect of rs73185306-T on the canonical function of piRNAs, namely, silencing TEs. Using data from the Genotype-Tissue Expression (GTEx) project, in which the frequency of rs73185306-T is 7.1%, we observed no effect of this SNP on expression of any human TE family (Figure S8). This is in contrast to the marked effect of inactivating *MOV10L1* mutations on TE expression in other mammals {Newkirk, 2017}. Therefore, the impact of this SNP on piRNA biology requires further investigation.

## DISCUSSION

Large WGS datasets offer a unique opportunity to study the human virome and human-virus coevolution. This is especially true in the case of integrated HHV-6, the result of unconventional virus-to-host horizontal gene transfer that violates “Biology’s Second Law” - Weismann’s proposed barrier preventing gene flow from the soma to the germline {Mattick, 2012}. However, even the prevalence of this interesting condition remains uncertain. Others have discussed that the commonly-cited prevalence 1% in the global population is an overestimate of the prevalence in healthy subjects, perhaps driven by inclusion of studies analyzing patient samples, among which the prevalence appears to around 2% {Pellett, 2012}. In that context, it is notable that less than 1% of the subjects in the diverse populations, some healthy (e.g. 1kGP), and others disease-enriched (e.g. BBJ, MSSNG), screened for integrated HHV-6 using WGS. Our observation that some subjects retain only a portion of the integrated viral genome, a single DR region, has not been reported by previous WGS-based screens. Depending on the region(s) of the viral genome targeted, the single DR form may have been detected in screens using other methods; if so, it was considered together with full-length HHV-6 integration. While this form is not a potential source of viral reactivation *de novo* from the host genome, the DR region encodes genes as well as microRNAs which may influence host or exogenous viral gene expression {Tuddenham, 2012}. Considering the existence of single DR integration as a distinct category of integrated HHV-6 is important for future studies on the implications of HHV-6 integration for human health.

In addition, the incidence of HHV-6 integration has remained unclear in the 20 years since this phenomenon was first described. However, careful studies using even small cohorts have been able to infer that ancestral integrations could account for much of integrated HHV-6 {Kawamura, 2017}. Recent work in Europeans, focusing on viral rather than human genetic diversity, has also suggested that integrated HHV-6 can reflect ancient, ancestral integrations {Zhang, 2017}. Our analysis of integrated HHV-6 in Japan clearly shows that integration of HHV-6 most often reflects ancestral, rather than incident, events in this population. We also defined the upper limit of the incidence of integration of HHV-6 in North America using WGS data from families. Because we did not observe any new integrations, larger family studies and phylogenetic analyses are needed to quantify this further. Our work clarifies that the evolution of integrated HHV-6, which may be more precisely described as “endogenous HHV-6” in examples for which stable germline inheritance is demonstrated, influences the interpretation of any associated human chromosomal SNPs {Liu, 2018}. Considering the incidence and prevalence of chromosomally-integrated forms of HHV-6 is important for properly interpreting the association of HHV-6 with phenotype and disease {Gravel, 2015}{Readhead, 2018}{Dowd, 2017}.

By providing molecular resolution of the relatively common East Asian endogenous HHV-6A integration breakpoint, our work advances the study of HHV-6 from a human genetic and paleovirological perspective. Unexpectedly, our study demonstrates that the majority of integrated HHV-6 in the Japanese population is located at the same cytogenetic locus, with independent integrations of both HHV-6A and HHV-6B present in the telomere of chr22q. Further study of this phenomenon is needed. The hypothesis that polymorphisms of this subtelomere prevent piRNAs from blocking HHV-6 integration, as they have been shown to do for other mobile genetic elements that can invade the germline, was a provocative one. In known examples of piRNA-guided silencing of TEs, integration of the element into a genomic locus that produces piRNAs is required {Duc, 2019}. In some species, the telomeres are in fact a piRNA-generating locus. For example, piRNAs are produced from telomere-integrated retrotransposons in flies, silkworms as well as the large TEs known as “terminons” in the telomeres of rotifers {Arkhipova, 2017}. Small RNA molecules described as piRNA-like RNAs have been reported to derive from mouse telomeres {Cao, 2009}. Whether telomere-integrated HHV-6 can act as a template to produce piRNAs, potentially protecting the germline from subsequent HHV-6 integration, remains to be tested. However, our work suggests that the proposed mechanism to explain the association between *MOV10L1* and integrated HHV-6, i.e. that piRNAs usually block HHV-6 integration but do not do so efficiently in subjects with a SNP affecting *MOV10L1*, is not supported by a number of independent integration events attributable to this SNP. At least in Japan, there seems to have been only one such integration; this same endogenous virus also exists in China and likely arrived to Japan from continental Asia.

What then explains the relatively prevalent endogenous HHV-6 variants, from both HHV-6A and HHV-6B, in the telomere of chr22q in Japanese subjects? We cannot exclude stochasticity, i.e. that it reflects two independent founder effects. If we assume that such a founder effect would be equally likely to be observed for HHV-6 integrated into any chromosome arm, the likelihood of observing both on the same chromosome arm is low, approximately 1/46^2^. Integration into chr22q may be favorable for some other reason, for example, related to the chromatin state of this subtelomere in the nuclei of cells of the germline, in which inherited integrations must take place. Another possibility is that integration itself occurs into all chromosomes, but is more readily lost when integrated onto other chromosomes. piRNAs have previously been associated with human recombination hotspots {Camara, 2016}, and the PIWI domain of prokaryotic argonaute proteins directly influences recombination {Fu, 2019}. Perhaps the reported *MOV10L1* SNP remains linked to HHV-6A by influencing the rate of recombination between the integrated virus and itself, however this would only explain one of the independent integrations observed on this chromosome arm.

Chr22q has also been reported to carry the penultimate shortest human telomere, longer only than that of 17p {Martens, 1998}. Notably, 17p has also been shown to harbor multiple HHV-6 integration events in Europeans {Torelli, 1995}{Zhang, 2017}. Both of these chromosome arms are also associated with subtelomere deletion syndromes, chr22q13 deletion syndrome and 17p13 monosomy {Bonaglia, 2011}{Stratton, 1984}, which likely result from telomere shortening and total loss. Natural selection could feasibly preserve haplotypes of these subtelomeres that avoided such losses {Hemann, 2001}. The use of mobile DNA to extend linear chromosome ends and maintain stable chromosome replication has emerged many times during eukaryotic evolution {(Saint-Leandre, 2019}, reviewed in{Kordyukova, 2018}), and remains a strategy used by organisms normally dependent on telomerase when it is absent {Begnis, 2018}. Analyses of more populations are needed to address the possibility that human chromosome arms with short telomeres benefit from carrying endogenous HHV-6 {Koonin, 2018}.

We used two methods to estimate the timing of integration of the endogenous HHV-6A variant prevalent in East Asian populations. The estimates suggest that HHV-6A integrated into the chromosome of an ancestral continental East Asian and arrived later in Japan. This model is consistent with human phylogeography and the observed distribution of rare alleles present on this haplotype in other populations. However, the confidence intervals of the two estimates of arrival to Japan do not overlap. The estimate based on mutation accumulation, which has been used previously to provide reasonable estimates of HHV-6 integration timing, places the arrival of this allele to Japan in the more distant past than that based on recombination. Perhaps this HHV-6A variant is accumulating mutations more rapidly than expected for chromosomal sequences. However, while the sample size is small, the observed polymorphisms in the integrated HHV-6A sequence do not evidence selection for nonsynonymous and potentially virus-inactivating mutations. Another possibility is that the haplotype is recombining less frequently than expected. The latter could support that the HHV-6A-linked haplotype is evolving under positive selection in Japan, although an effect of the viral genome on homolog pairing and synapsis cannot be excluded. Our current study is underpowered to further address this intriguing possibility.

We described the molecular signature of HHV-6 excision, via recombination, from its position in telomere for the first time. As extensively characterized *in vitro*, this event likely represents viral reactivation, with potential production of infectious virus. We observed this form in nine subjects in BBJ, about 30% of all subjects with some form of integrated HHV-6B sequence. Our data are consistent with germline transmission of the single DR form. We thus suspect that these represent a single historical reactivation event in the germline, but nevertheless support that this process does occur *in vivo*, not only *in vitro*. Distinct variants of the integrated single DR form were also observed in MSSNG subjects. These data confirm that the risk of excision and thus potential reactivation of integrated HHV-6B is real, though how often this occurs remains difficult to quantify. Studies with somatic tissues sampled from many sites may be useful for this purpose {Peddu, 2019}. While recombination resulting in a single DR “scar” is one of the proposed routes of excision and reactivation, another involves “scarless” excision due to recombination of telomeric repeats flanking the virus genome{Wood, 2017}. WGS analysis is unable to infer this type of excision. Our results support caution in using cells and tissues from subjects bearing integrated HHV-6 for transplantation; upon immunosuppression, exposure to HHV-6 excised from donor cells may be harmful {Hill, 2017}{Bonnafous, 2018}. More generally, these results support the concept that HHV-6 excision from destabilized telomeric heterochromatin, for example in the aged, may contribute to human disease {Gravel, 2015}{Dowd, 2017}{Readhead, 2018}. With data from completed or ongoing population WGS projects, a global assessment of integrated HHV-6 prevalence, evolution, and association with disease should soon be possible. In addition, understanding any immune responses engendered by endogenous HHV-6, either conventional {Peddu, 2019} or genomic {Ophinni, 2019}, are relevant to understanding the biological significance of this phenomenon.

## METHODS

### Screening of HHV-6 carriers based on WGS

A total of 7,485 WGS samples were obtained from the BBJ project {Hirata, 2017}{Nagai, 2017}. Read alignment to the human reference genome hs37d5 and variant calling were previously performed for 3,256 high-depth WGS samples as described elsewhere {Okada, 2018}. Additionally, 4,229 low-depth WGS were analyzed with Genomes on the Cloud (GotCloud) pipeline{Jun, 2015}. From each BAM file, we extracted unmapped reads. We required that both paired-end reads were unmapped. We realigned unmapped reads to the HHV-6A genome (KJ123690.1) using the BWA-MEM algorithm (BWA version: 0.7.13) {Li, 2009}. Read depth was measured as the mode of per-base read depth across the length of the viral reference genome. The threshold of 30% depth relative to WGS depth of coverage was chosen in order to capture those with inconsistent mapping or some degree of acquired somatic mosaicism, but exclude those with viremia, which has resulted in 10-1000x lower coverage in other studies {Moustafa, 2017}{Liu, 2018}.

### Kinship Analysis or Genetic correlation matrix

We evaluated genetic relatedness between subjects based on genotypes of common variants across the genome by plink software {Purcell, 2007}. We excluded the HLA region and restricted variants with minor allele frequency more than 5% and not in linkage disequilibrium with other variants (r^2^ > 0.2). PI_HAT, the proportion of identity by descent (IBD) defined as probability (IBD=2)+0.5×probability (IBD=1), was computed to determine genetic relatedness (PI_HAT> 0.25).

### GWAS of HHV-6

We performed variant joint-calling for high-depth WGS (N = 3,262) by aggregating individual gVCF with GATK following the current recommended best practice. Briefly, variant QC was conducted for samples in two subsets 1) samples sequenced at 30X (N = 1,292), variants that meet any of the following criteria (1) DP < 5, (2) GQ < 20, or (3) DP > 60, and GQ < 95 were removed; 2) samples sequenced at 15X (N = 1,964), variants that meet any of the following criteria (1) DP < 2, (2) GQ < 20 were removed. We phased the resulting genomes for use as a reference panel using SHAPEIT2 and imputed the variants of low-depth WGS using IMPUTE2{Howie, 2012}. The final variant set include both high-depth and low-depth WGS. We performed exact test to compare the allele frequency between integrated HHV-6 and non-carriers for all variants using Plink (version 1.9). 5 × 10−8 is used as threshold to define the genome-wide significance levels.

### Viral phylogenetic analysis

We reconstructed the HHV-6 viral genome of 10 subjects with high HHV-6 read depth who have been sequenced at high-depth (4 HHV-6A and 6 HHV-6B). First, reads mapped to HHV-6 were further aligned against the integrated HHV-6A genome derived from a Japanese individual NA18999 (GenBank Accession number: KY316047.1) using BWA. Based on the alignment, variants calling was performed using freebayes (version: v1.2.0-2-g29c4002) with parameters ploidy = 1 and min-alternate-fraction = 0.8 {Garrison, 2012}. Resulting variants were patched into the reference genome to obtain the sample-specific iciHHV-6 viral genome. These 10 samples, together with previously reported chromosomally-integrated or nonintegrated HHV-6 genomes from the literature (N=19), were used for viral phylogenetic analysis. We extracted and concatenated sequences of 3 viral genes (U27, U43, U83) from each sample. Trees were built using the neighbor-joining algorithm with 1,000 times bootstrap using MEGA7 software{Kumar, 2008}. The HHV-6 genomes used in this analysis and their GenBank Accession numbers are shown in Table S7. We used the same method for phylogenetic analysis of integrated HHV-6B and others by concatenating shared variant sites that were called in all subjects.

### Haplotype phasing for variants in chr22q region

We extracted 174 variants within chr22q sub-telomeric region for high HHV-6 read-depth individuals (N = 32) and unrelated HHV-6-negative subjects (N = 100) from the microarray dataset of BBJ as described previously {Kanai, 2018}. Genotype data of two subjects with integrated HHV-6 for whom sequence data was available through the 1000 genome project (NA18999 and HG00657) were also added {Genomes Project, 2015}. A telomeric variant was appended to reflect presence or absence of integrated HHV-6 based on WGS. We used PHASE to infer the individual haplotypes, and subsequently generated the phylogenic tree based on neighbor-joining (NJ) method with MEGA (version 7) {Stephens, 2003}.

### Sanger sequencing of subjects with low-depth WGS or FISH-mapped integrated HHV-6

Sanger sequencing was conducted to genotype four variant sites including rs73185306, rs149078280, rs566665421 and chr22_51184036_C_G for BBJ subjects with integrated HHV-6 sequenced by low-depth WGS and an additional 9 Japanese subjects with integrated HHV-6 previously mapped by FISH. The PCR primers and sequencing primer sequences are provided in Table S8. Briefly, we used 10 ng of genomic DNA for PCR amplification. Purified PCR products were sequenced on an Applied Biosystems Genetic Analyzer 3130 (Thermo Fisher Scientific, MA, USA) with BigDye Terminator v3.1.

### Estimation of the age of integrated HHV-6A

The first method to estimate the timing of the shared HHV-6A integration is based on the assumption that the observed Chinese and Japanese integrated HHV-6A genomes derived from a single integration event, and after being integrated the mutation rate of the integrated HHV-6A genome is the same as that of other human chromosomal DNA. By considering variants unique to one or more variants but absent in the consensus of all variants (interpreted to be *de novo* mutations arising after integration), we estimated the expected age and 95% CI according to the Poisson distribution. The human mutation rate is estimated at 1.2*10^-8 per site per generation and we assume 25 years between generations {Kong, 2012}. We excluded the repeat-rich, 2-copy DR regions and considered only variants arising in a 140 kb unique (U) genic region. We performed joint calling using FreeBayes for 4 BBJ integrated HHV-6A and HG00657 and NA18999, and subsequently filtered out unique mutations and visually confirmed the mutations by IGV.

The second method is based on the decay of linkage disequilibrium (LD) between iciHHV-6A and a linked marker allele (rs149078280-T). The population frequencies of iciHHV-6A allele, rs149078280-T allele, and haplotype bearing these two alleles are denoted by *p*, *q*, and *h*. The recombination rate between two loci is denoted by *c*. In this setting, *h* in the next generation is given by

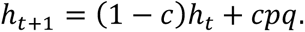

The extent of LD between the two loci (*δ*) is characterized by the fraction of rs149078280-T allele among integrated HHV-6A-bearing chromosomes (i.e., *h*/*p*). Dividing the above equation by *p*, we obtain

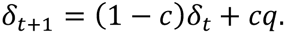

At the time of integration (*t*=0), *δ* is 1 and then decreased exponentially at a rate *c* to *q* as generation passes. Since integrated HHV-6A is rare, the hitchhiking effect of integrated HHV-6A on the frequency of rs149078280-T (*q*) is very limited. It is therefore assumed that *q* has been constant since the integration. Solving the above recurrence equation, the current *δ* is given by

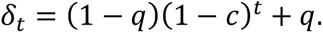

Solving this for *t*, we obtain

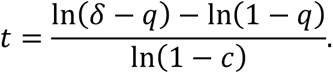

The observed values of *δ* and *q* were 0.666 (=8/12) and 0.000313. In this study the position of rs149078280 on chromosome 22 (73.29267 cM) was retrieved from genetic maps for the 1000 Genomes Project variants (https://github.com/joepickrell/1000-genomes-genetic-maps). The integrated HHV-6A genome is located telomeric to the terminal nucleotide present in the reference sequence of chr22q. Therefore we used a value of 74.10956 cM, which represents the distance from rs149078280 to the most telomeric informative allele on this chromosome. This value derives from subtelomeric markers, not the telomere itself; the genetic distance is thus a conservative underestimation of the actual distance. Accordingly, the recombination rate (*c*) between integrated HHV-6A and rs149078280 was assumed to be 0.00817 (i.e., 0.01x[74.10956-73.29267]). Putting *δ*, *q*, and *c* into the above equation, *t* is 35 (generations). If we assume the human generation time is 25 years, this corresponds to approximately 875 years ago. The estimated age is dependent on the numbers of haplotypes observed (i.e., observed haplotype frequencies). The haplotype frequencies used here are estimated based on a random set of 14,970 chromosomes from Japanese subjects. The numbers of integrated HHV-6A _ rs149078280-T, integrated HHV-6A _ rs149078280-C, without integrated HHV-6A _ rs149078280-T, and without integrated HHV-6A _ rs149078280-C haplotypes were 9, 3, 6, and 14,952 respectively. To obtain an empirical bootstrap confidence interval, we generated 100,000 bootstrap samples, each of size 14,970, that satisfied the condition that four different haplotypes were observed and *δ*-*q*>0. The 95% confidence interval of age was 10-94 generations (250-2,350 years) (Figure S2B).

### Nanopore long read sequencing

We obtained lymphoblastoid cell lines (LCLs) of HG00657 and NA1899 from the Coriell Cell Repositories and cultured them according to the protocol provided. We extracted high molecular weight (HMW) DNA using Gentra Puregene Cell Kit (Qiagen, Hilden, Germany) from 5 × 10^6^ cultured cells. DNA was quantified with a Qubit fluorometer (Invitrogen, US) and 1 ug of HMW DNA was used to construct the DNA library using Nanopore ligation sequencing kit SQK-LSK109 (Oxford Nanopore Technologies, ONT, Oxford, UK). We loaded the library into R9.4 flow cell (ONT) and subsequently conducted sequencing on a MinION (ONT) sequencer. Base calling for the MinION raw sequencing data was done on a MinIT (ONT) via MinKNOW software (ONT). We used minimap2 to align the reads against a customized hg19 reference genome in which the HHV-6A reference genome has been added as a decoy sequence. The inferred integration breakpoint is available upon request and will be deposited to the DNA Data Bank of Japan upon publication.

### Integrated HHV-6 in the MSSNG database

MSSNG database (https://www.mss.ng) release 5 was accessed via the Google Cloud Genomics platform. A total of 5,555 subject’s WGS datasets were available at the time of access. We screened for subjects with integrated HHV-6 using pre-available unmapped bam files using the same approach described above to screen BBJ subjects. This analysis was performed using Google Cloud Genomics platform. We queried the genotype of rs73185306 from the MSSNG dataset using Google BigQuery. We tested for an association between ASD and HHV-6 integration using transmission disequilibrium test (TDT) {Spielman, 1994}. We tested for Mendelian error by determining if any parents without integrated HHV-6 gave rise to children with integrated HHV-6, which would indicate newly incident HHV-6 integration. The rule of three was calculated as 3/1766 {Hanley, 1983} and the upper limit of the Wilson score interval was calculated as described previously {Wilson, 1927}.

### Digital droplet PCR (ddPCR)

We conducted ddPCR to determine the existence of DR and U region of HHV-6 and to distinguish the 6A/6B subtype for subjects with low depth of coverage of HHV-6 (N = 9), and control subjects carrying integrated HHV-6A (N = 2) or HHV-6B (N = 2) from BBJ. We used a primer/probe set for HHV-6 U57 gene and RPP30 (autosomal control) as previously described {Sedlak, 2014}. For the DR region, we designed common primers and species-specific probe for 6A and 6B by identifying a region of species-specific variation using an alignment of reference HHV-6 sequences (Figure S1) and those identified in our study (Figure S. Primer and probe information is provided in Table S9. To prepare the ddPCR reaction mix, 10 µl of 2× ddPCR Supermix for Probes (Bio-Rad, Hercules, CA), 1 µl of each 20× primer-probe mix (18µM each PCR primer, 5µM probe), and 15 ng genomic DNA in a final volume of 20µl. The reaction mixture was loaded onto a DG8 cartridge (Bio-Rad) with 70µl of droplet generation oil (Bio-Rad) and processed in the Droplet Generator (Bio-Rad). After droplet generation, 40 ul droplets were transferred into a 96-well plate and proceed to thermal cycling with the following conditions: 95°C for 10 minutes, 94°C for 30 seconds and 60°C for 1 minute for 40 cycles and ending at 98°C for 10 minutes. After amplification, the droplets were read by the Droplet Reader (Bio-Rad). QuantaSoft analysis software (V1.3.2.0) was used for data analysis and quantified copy number of target per µl was obtained and analyzed.

### Transposable element expression analysis

Use of GTEx data was authorized by NHGRI Data Access Committee via dbGaP (https://www.ncbi.nlm.nih.gov/gap/) (Project: 19481). RNA-Seq data of 163 testis samples in GTEx were downloaded from SRA (https://www.ncbi.nlm.nih.gov/sra) using SRA Toolkit (https://www.ncbi.nlm.nih.gov/sra/docs/toolkitsoft/). After read trimming by Trimmomatic (0.36) {Bolger, 2014}, reads were aligned to the human reference genome (GRCh38.p12) using STAR (2.6.0) {Dobin, 2013} with the annotations of genes and TEs. As sources of the annotations of genes and TEs, GENCODE22 (https://www.gencodegenes.org/) and RepeatMasker at UCSC (http://genome.ucsc.edu/) were respectively used. Read count matrix was generated using featureCounts (v1.6.3) {Liao, 2014}, and the expression levels of genes and TEs were normalized using the variance-stabilizing transformation function implemented in DESeq2 {Love, 2014}. Genotype information of the subjects was obtained from dbGaP.

### Data availability

WGS data of a part of the BBJ subjects (n = 1,026) is publicly available at the National Bioscience Database Center (NBDC) Human Database (https://humandbs.biosciencedbc.jp/en/) under the research ID hum0014, and remaining data are available on request after approval of the ethical committee of RIKEN Yokohama Institute and the Institute of Medical Science.

## Acknowledgements

We appreciate the staff of BBJ for their excellent assistance in collecting samples and clinical information. We thank Dr. Stephen S Francis (University of California, San Francisco), Dr. Amr Aswad (Free University of Berlin), Dr. Ruth F Jarrett (University of Glasgow), Dr. Wayne Yokoyama (Washington University in St. Louis) and many other participants in the 11^th^ International Conference on Human Herpesvirus 6 and 7 for helpful discussions. We thank Dr. Youdiil Ophinni (Kobe University) for assistance with figures. The authors wish to acknowledge the resources of MSSNG (www.mss.ng), Autism Speaks and The Centre for Applied Genomics at The Hospital for Sick Children, Toronto, Canada. We also thank the participating families for their time and contributions to this database, as well as the generosity of the donors who supported this program. The Genotype-Tissue Expression (GTEx) Project was supported by the Common Fund of the Office of the Director of the National Institutes of Health, and by NCI, NHGRI, NHLBI, NIDA, NIMH, and NINDS. The data used for the analyses described in this manuscript were obtained from: dbGaP accession number phs000424.v7.p2 on 4/23/2019.

---

**Figure S1.**
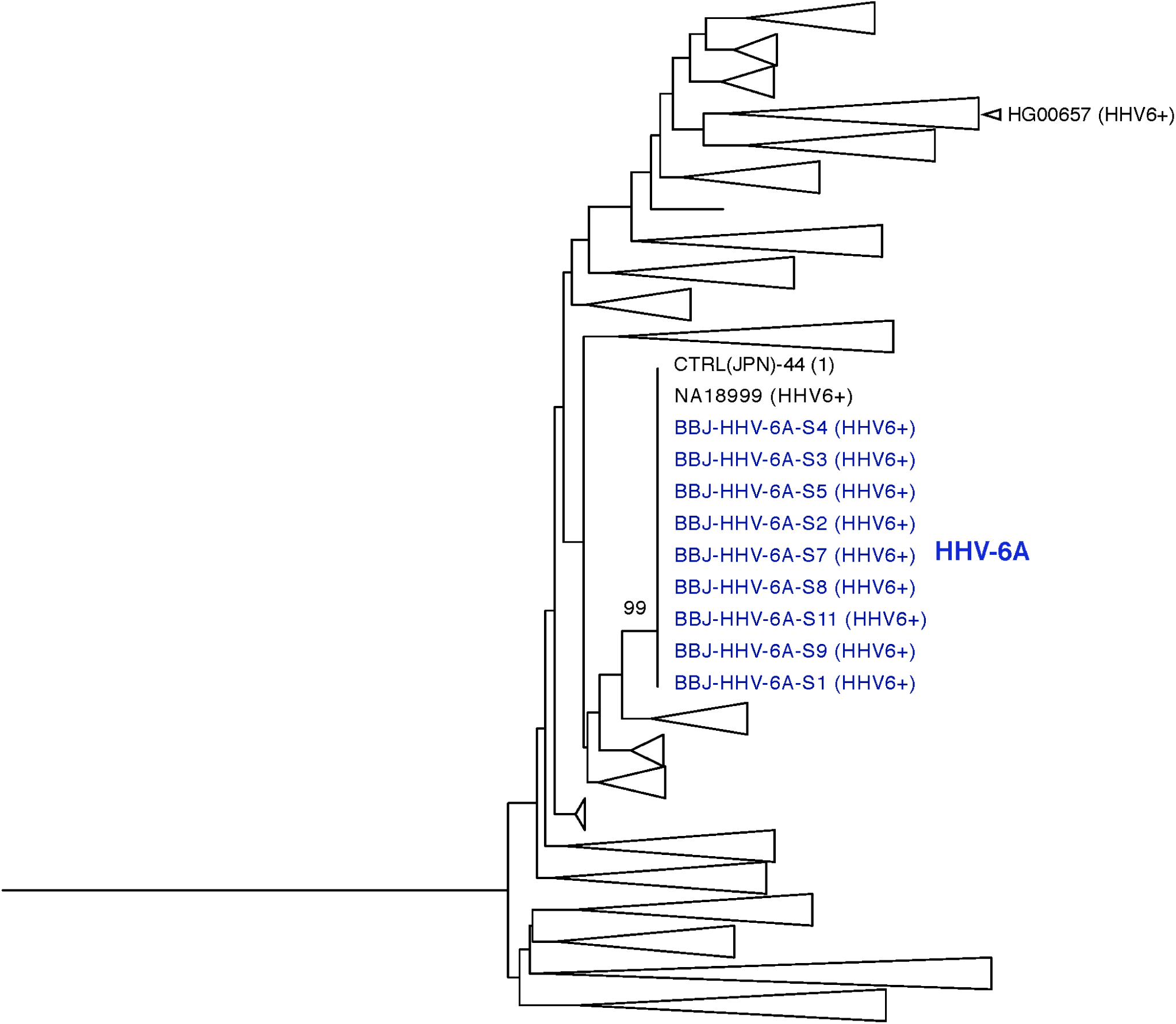
Neighbor-joining phylogenetic tree of phased 22q subtelomeric haplotypes with HHV-6A cluster highlighted. SNPs of 22q subtelomere were phased to obtain estimates of individual haplotypes for 32 subjects with high HHV-6-mapping read depth, 100 control subjects without HHV-6-mapping reads, and subjects NA18999 and HG00657 (data from 1kGP). Branches containing the clustered HHV-6A-associated haplotype is shown expanded (see figure S2 for fully expanded tree). Chinese subject HG00657 is highlighted with a blue triangle. The HHV-6 sequence carried by this individual is shared with BBJ HHV-6A subjects (Figure 3), yet the shared 22q subtelomeric haplotype is lost except for the most telomeric rare variant (Table 1) Bootstrap value per 100 replicates of selected nodes is shown. 174 SNPs were phased.

**Figure S2.**
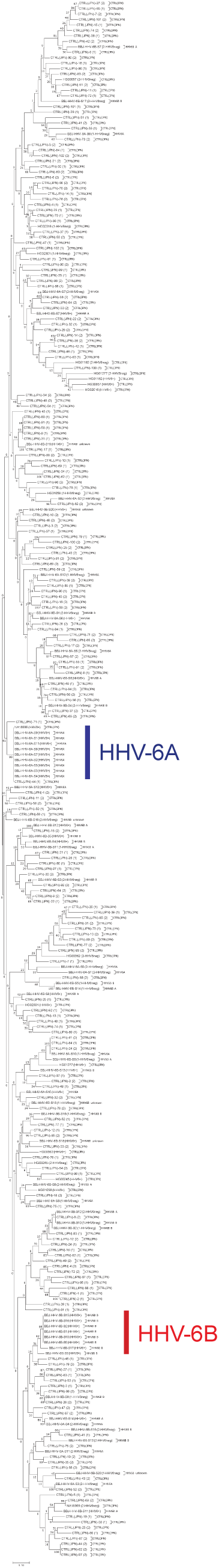
Neighbor-joining phylogenetic tree of phased 22q subtelomeric haplotypes from BBJ. SNPs of 22q subtelomere were phased to obtain estimates of individual haplotypes for 32 subjects with high HHV-6-mapping read depth, 100 control subjects without HHV-6-mapping reads, and subjects NA18999 and HG00657 (data from 1kGP). Lines and labels mark haplotype sharing among subjects with integrated HHV-6. Branches of this tree are selectively collapsed in figures S1 and S4B.

**Figure S3.**
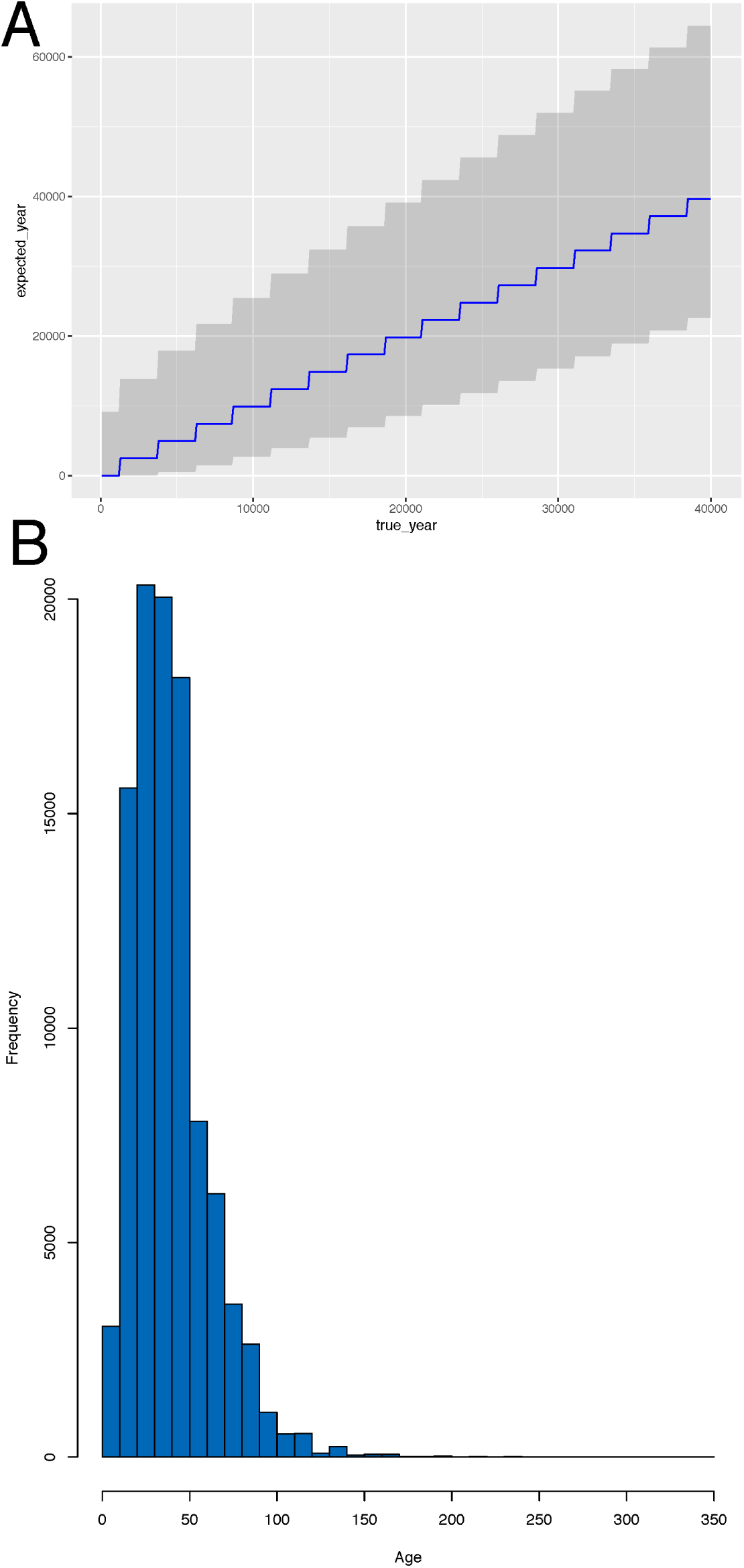
Estimated dating of East Asian endogenous HHV-6A integration. **A) Simulated age of integrated HHV-6A in Japan based on accumulated mutations.** We estimated the integration age by assuming that the mutation rate of the integrated HHV-6 genome is same as other human chromosomal sequences and each generation is 25 years. The blue line simulates the expected accumulation of mutations over time, the X axis indicates the true age, Y axis indicates the expected age calculated based on number of observed mutations, and the gray area represents the 95% CI of the expected age. **B) Empirical distribution of the age of integrated HHV-6A in Japan based on recombination.** Histogram showing the distribution of predicted age in generations (x-axis) of the iciHHV-6A allele obtained by 100,000 bootstrap samples.

**Figure S4.**
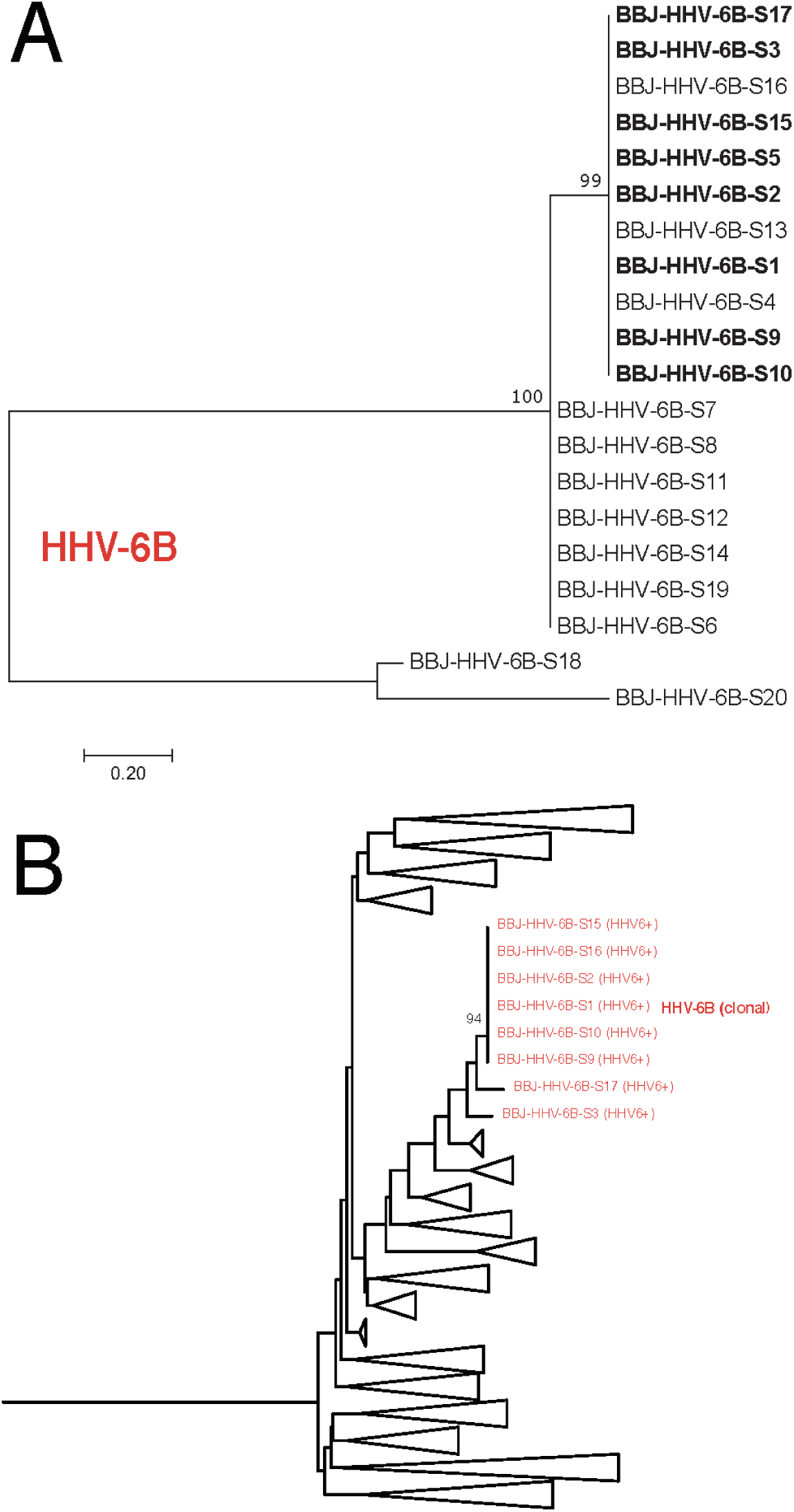
A clonal endogenous HHV-6B variant is present in Japanese subjects with a shared chr22q haplotype. **A) Clonal integrated HHV-6B evidenced by phylogenetic analysis.** Joint-calling of variants was performed for BBJ subjects with integrated HHV-6B of both high and low depth (N =20). 44 variant sites which were called in all subjects were selected and concatenated for phylogenic tree analysis using the maximum likelihood method. Subjects clustering by chr22q haplotype analysis, shown below, are bolded. Bootstrap value per 100 replicates of selected nodes is shown. The scale bar represents 0.20 substitutions per site. **B) Neighbor-joining phylogenetic tree of phased 22q subtelomeric haplotypes with clonal HHV-6B cluster highlighted.** SNPs of 22q subtelomere were phased to obtain estimates of individual haplotypes for 32 subjects with high HHV-6-mapping read depth, 100 control subjects without HHV-6-mapping reads, and subjects NA18999 and HG00657 (data from 1kGP). Branches containing clustered clonal HHV-6B associated haplotypes are shown expanded; Bootstrap value per 100 replicates of selected nodes is shown. 174 SNPs were phased.

**Figure S5.**
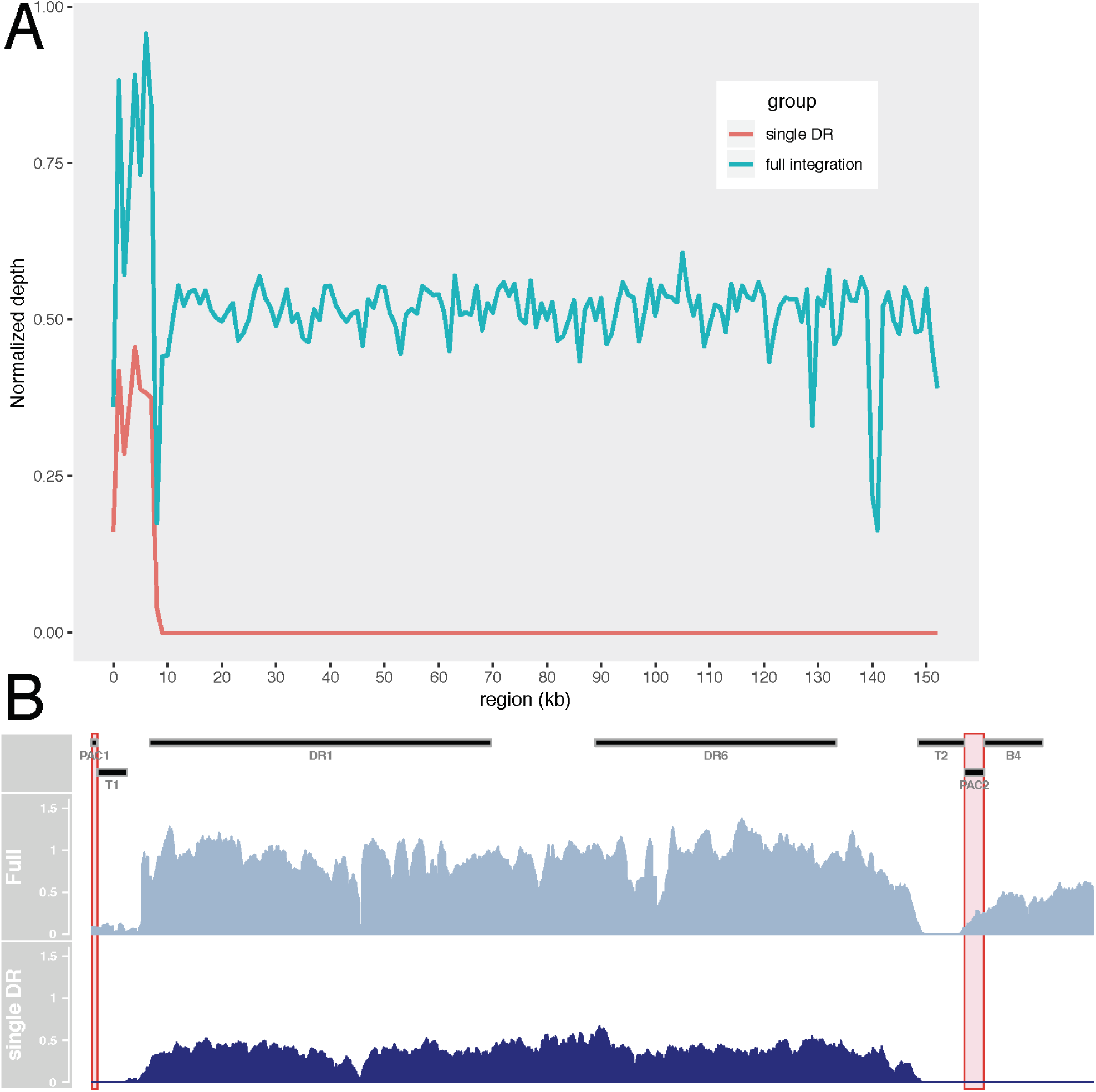
A subset of subjects with integration of a single HHV-6B DR. **A) Depth in subjects with reads mapping only to the DR region is half that of those with reads mapping across the entire viral genome.** We summarized the depth of coverage in 1 kb sliding window across the HHV-6B genome (x axis), the read depth from subjects in one of two groups is plotted (y axis). The average depth of subjects meeting the described threshold to infer integrated HHV-6 from high-depth WGS (N=10; 4 integrated HHV-6A and 6 HHV-6B) are shown in blue. The average depth of subjects with HHV-6-mapping reads below the threshold from high-depth WGS are shown in red (N = 4). Subjects failing to reach the threshold to infer integration of intact HHV-6 also produce reads of depth consistent with germline chromosomal integration of a portion of the HHV-6 genome, the DR region. A decoy HHV-6B reference genome with the DR(R) removed (which is identical in sequence to DR(L)) was used for mapping and calculation. **B) Zoomed view of coverage of the DR region.** Comparison of the depth of reads mapping to the DR region between those with full and single DR integration suggests that a single copy of the DR region remains in the latter. Pac-1 and pac-2 sequences important for viral genome packaging are highlighted in red. T1 and T2 are telomere repeat sequences. DR1 and DR6 are spliced open reading frames present in viral genome annotation ID #NC_000898.

**Figure S6.**
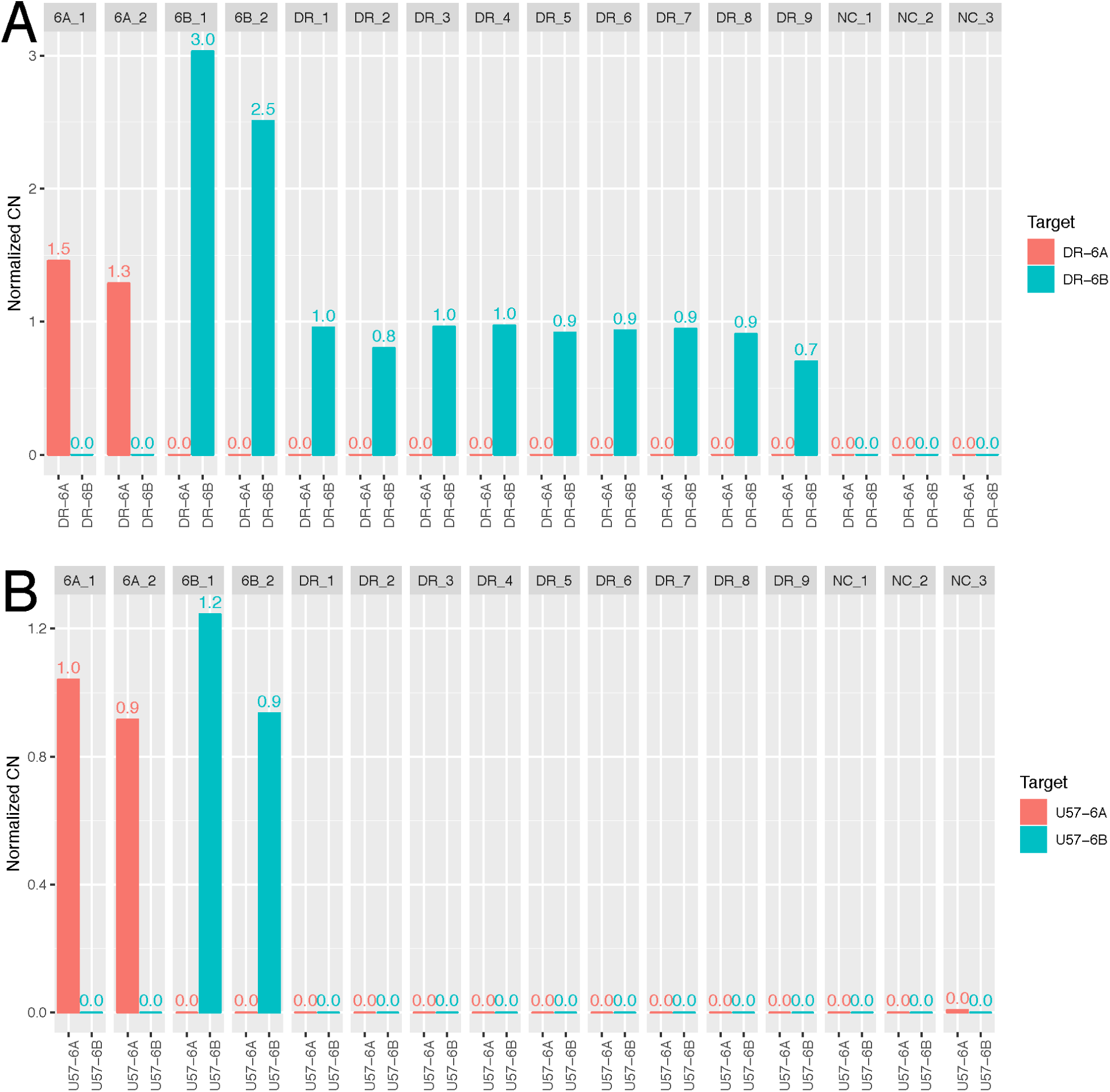
Droplet PCR to estimate the copy number of DR and U regions. **A) DR copy number normalized by RPP30.** The bar plot shows the normalized copy number (CN) for each individual determined by 6A/6B specific DR probe. NC, negative control. **B) U57 copy number normalized by RPP30.** The bar plot shows the normalized copy number (CN) for each individual determined by 6A/6B specific U57 probe. NC, negative control.

**Figure S7.**
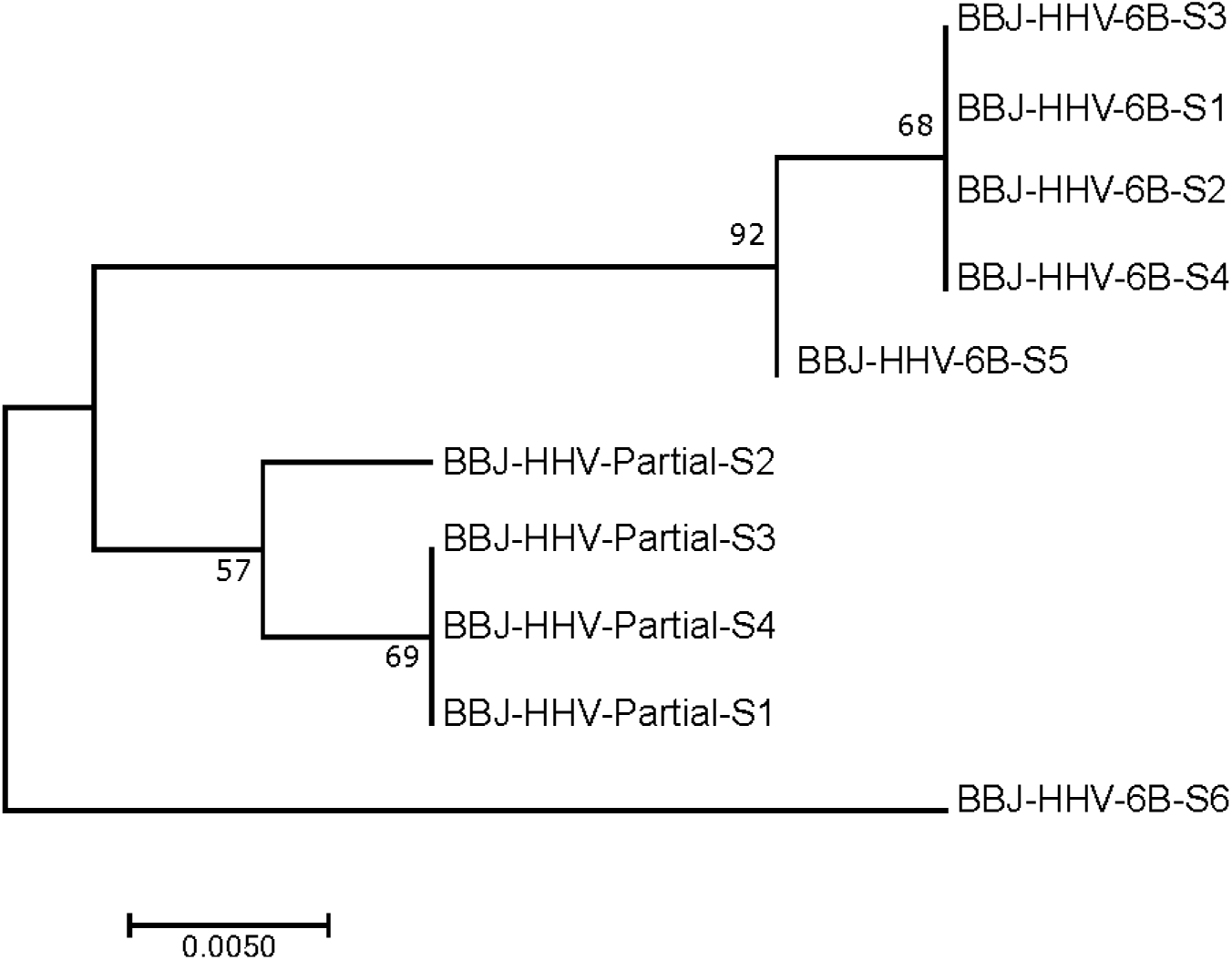
Phylogenetic analysis of integrated HHV-6B based on DR region. We performed joint-calling of variants in the DR region for BBJ subjects. We considered subjects sequenced at high-depth (N = 10) to exclude the possibility of inaccurately-called variants in the repeat-rich DR region in subjects with low read depth. A total of 237 variant sites in DR region which were called in all subjects were selected and concatenated to generate a phylogenic tree using the maximum liklihood method. Bootstrap value per 100 replicates of selected nodes is shown. The scale bar represents 0.005 substitutions per site.

**Figure S8.**
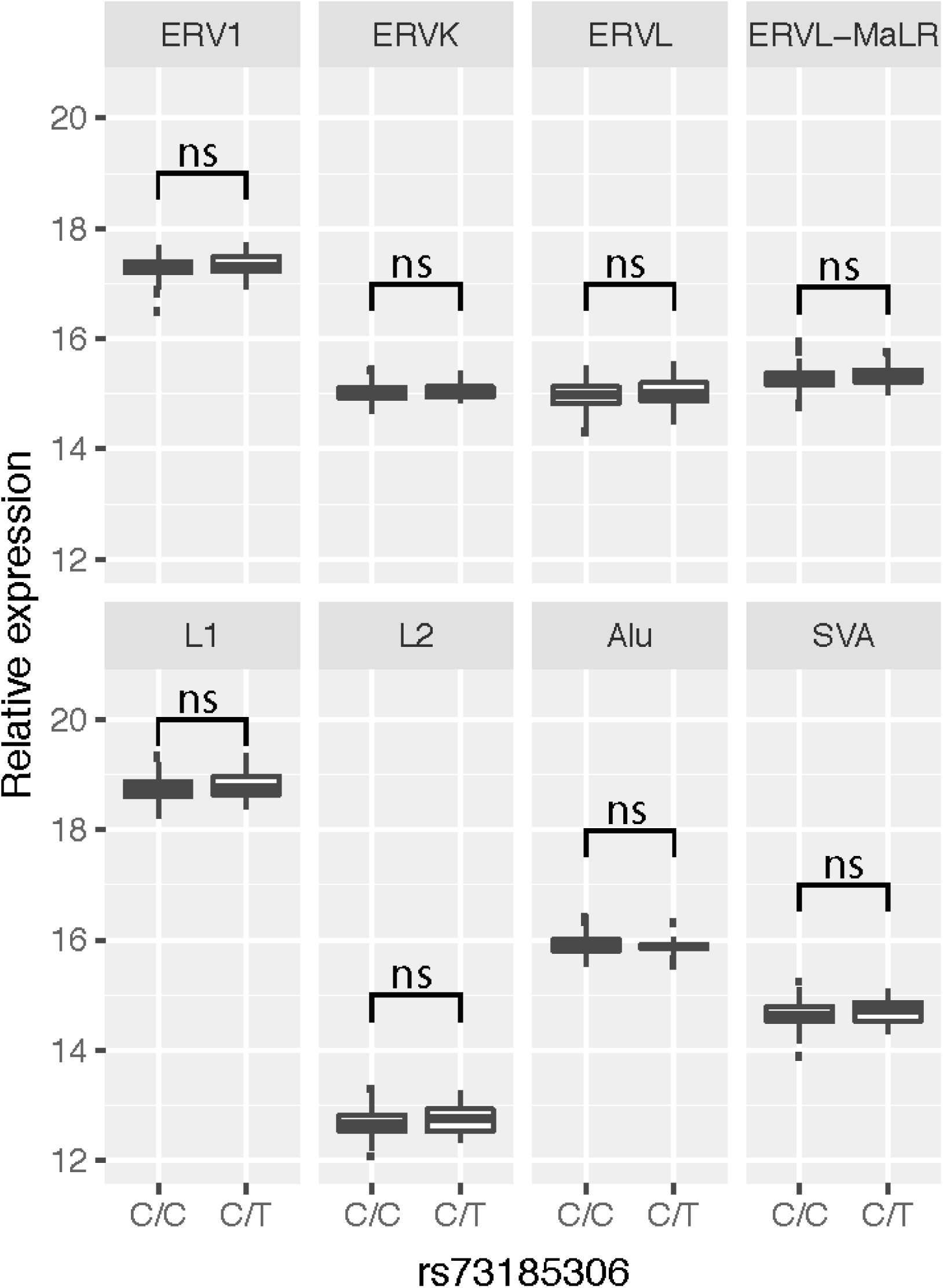
**rs73185306 is not associated differential RNA expression of transposable elements (TEs) in testis**. Testis RNA expression levels of various families of TEs are shown, as labelled, grouped by genotype at the index SNP reported by Liu et al. (n = 163; C/C = 143, C/T = 20).

Table S1: Genome-wide significant variants from GWAS of integrated HHV-6A/B

Table S2: Genome-wide significant variants from GWAS of integrated HHV-6A

Table S3: Sanger sequencing of additional Japanese subjects with integrated HHV-6A mapped by FISH to chromosome 22q

Table S4: Genome-wide significant variants from GWAS of clonal integrated HHV-6B

Table S5: Sanger sequencing of additional Japanese subjects with integrated HHV-6B mapped by FISH to chromosome 22q

Table S6: Association between rs73185306 and integrated HHV-6 in MSSNG database

Table S7: Additional HHV-6 genome sequences included in current study

Table S8: Primers used for Sanger sequencing

Table S9: Primers and probes used for digital droplet PCR

Supplemental Tables available upon request to

